# Structure of a pH-sensitive pentameric ligand-gated ion channel from the *Sarcoptes* scabies mite

**DOI:** 10.1101/2025.06.11.659021

**Authors:** Jessica Kleiz-Ferreira, Marijke Brams, Peter J. Harrison, Casey I. Gallagher, Mieke Nys, Ysaline Donze, Andrew Quigley, Daniel Bertrand, Chris Ulens

## Abstract

Scabies is a skin infestation caused by the *Sarcoptes scabiei* mite. It produces a substantial global health burden, which is exacerbated by emerging resistance to prominent treatments, such as ivermectin. An anionic pentameric ligand-gated ion channel (pLGIC) identified in the mite, termed as SsCl, shows unique pH-sensitivity and is significantly modulated by ivermectin. Here, we use cryo-EM and electrophysiology to explore the pH-sensing mechanisms of SsCl and the impact of ivermectin on channel activity. Structures of SsCl were resolved under various pH conditions to capture a closed (pH 6.5) and desensitized (pH 9) state, alongside ivermectin-bound conformations. The desensitized structure adopts an unexpected “hourglass” conformation, suggesting the gating mechanisms may be more related to cation-selective channels. Structure-based analysis and mutagenesis highlighted extracellular histidine and glutamate residues which act as pH-sensors, revealing the protonation-dependent gating mechanism. Ivermectin binds at transmembrane subunit interfaces, stabilizing an open-pore conformation through hydrophobic interactions. Ivermectin bound-structures reveal a pH-dependent modulation, enhancing open-state prevalence at pH 9 and enabling atypical activation at pH 6.5, consistent with electrophysiological data. These structural and functional insights elucidate SsCl’s unique pH sensitivity and ivermectin’s mode of action, providing a foundation for designing next-generation therapeutics targeting this pathogen-related ion channel.

## Introduction

pH-sensitive chloride channels (pHCls) are unique members of the pentameric ligand-gated ion channel (pLGIC) family (for review, see^1^) which are primarily found in invertebrates like insects^2^ and arachnids^3^. These channels are specifically activated by changes in pH, which occurs during metabolic shifts or stress responses^4–6^. When activated by alkaline pH, they allow the passage of chloride ions through the cell membrane which helps regulate cellular homeostasis, maintain ionic balance and mediate signal transduction^4,5,7^. Unlike traditional ligand-gated receptors which have well-characterized binding cavities, the agonist-like region involved in the pH-sensing and its associated gating mechanisms are not well known. Further research on pHCl channels would not only contribute to our understanding of ion channel physiology but also holds promise for the development of targeted treatments against parasitic infections and agricultural pests, due to their selective expression in invertebrates^8^.

One of the most medically relevant parasitic infestations is scabies, a highly contagious ectoparasitic skin condition caused by *Sarcoptes scabiei* (itch mite)^9^. This tiny mite penetrates the upper layer of the skin, where it lays eggs and induces an immune response that leads to intense itching, inflammation, and characteristic skin rashes. The infestation often causes severe discomfort and can lead to secondary complications, such as impetigo, rheumatic fever and chronic kidney diseases^10,11^, if left untreated. Scabies is estimated to affect 200-300 million people worldwide, particularly in impoverished communities or crowded environments, such as nursing homes, schools, and dormitories^9,12^. The World Health Organization has therefore highlighted scabies as a disease of public importance and lists it as a neglected tropical disease^13,14^.

The standard treatment for scabies involves topical medications, such as permethrin cream, or oral medications like Ivermectin (IVM)^15^. IVM is a broad-spectrum antiparasitic medication originally developed in the late 1970s^16,17^. Due to its high efficacy and low toxicity, it has since become a cornerstone in the treatment of parasitic conditions in both humans and animals, and was recognized with the 2015 Nobel Prize in Physiology or Medicine^18^. Its use in mass drug administration campaigns has significantly reduced the prevalence of parasitic diseases and improved public health in endemic regions^19^. However, the emergence of drug resistance in some parasitic populations^20^ highlights the need for ongoing research into the activity of IVM at pharmacologically relevant targets, while stimulating the development of new treatments.

IVM binds to chloride channels in the nerve and muscle cells of parasites, causing paralysis and death of the parasite while having minimal effects on human cells^21^. Its activity was originally characterized using the invertebrate glutamate-gated chloride channel GluCl^22^, but has since been shown to act on other pLGICs including GABA receptors^23^, glycine receptors^24^ and insect pHCl channels^25^. However, its exact mechanism of action in *Sarcoptes scabiei* remains unknown.

One likely target is a pHCl channel identified in *Sarcoptes scabiei,* SsCl. This channel is closely related to various glycine and pHCl channels from insects (Sup.Fig. 1 and 2). In particular SecCl channels, which were originally identified in *Drosophila* and have been shown to mediate hormone-induced fluid secretion^25^. SsCl is activated by alkaline environments and modulated by IVM, which acts slowly and pseudo-irreversibly, even at acidic pHs when the channel is closed^2,3,7^. However, its functional role in *Sarcoptes scabiei* and how these mechanisms occur remain unknown. SsCl presents a novel target for future scabies treatments and an opportunity to explore the poorly understood mechanisms of pHCl channels.

**Figure 1.**
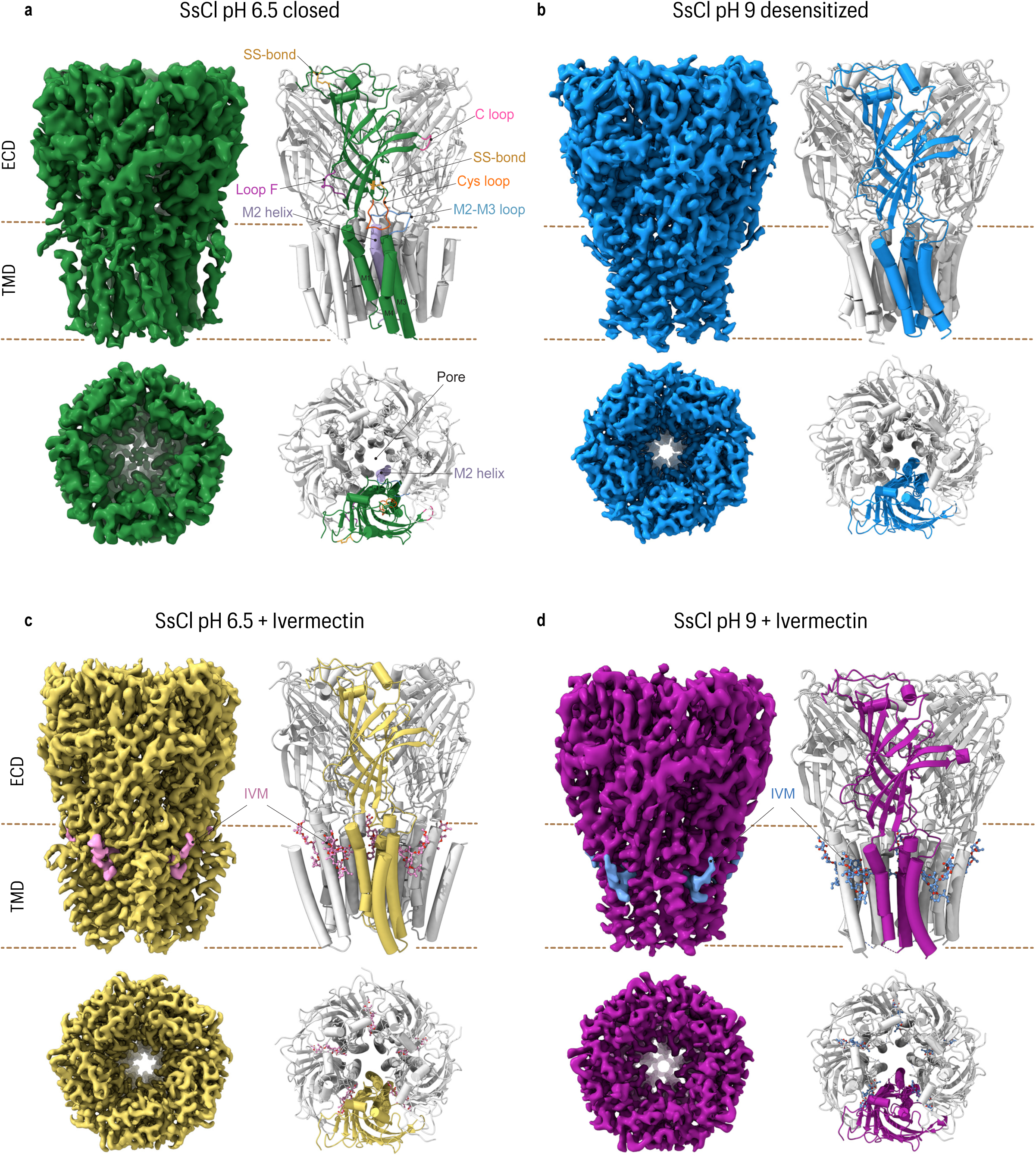
Cryo-EM structures of SsCl channel in four conformational states. Top (side view) and bottom (top view) panels show the cryo-EM density map (left) and atomic model (right) of SsCl under four conditions: **a**, pH 6.5 (closed state); **b**, pH 9.0 (desensitized state); **c**, pH 6.5 with IVM, shown in pink; and **d**, pH 9.0 with IVM shown in blue. In **a**, key structural elements, including loops and transmembrane helices, are annotated on the model.

**Figure 2.**
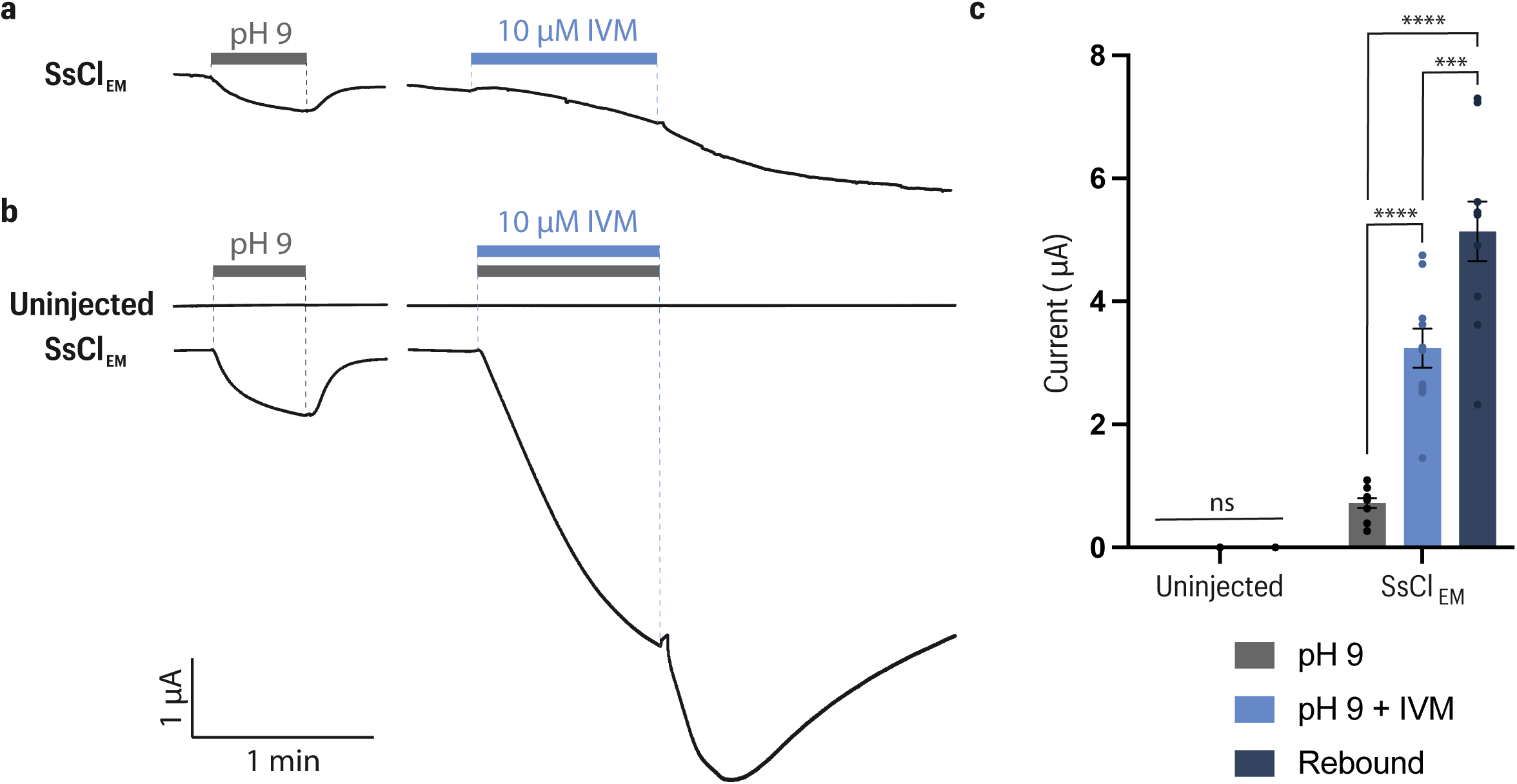
Effect of ivermectin on SsCl. **a**, Representative traces from oocytes expressing SsCl_EM_ illustrating an application of pH 9 (grey bar) followed by an application of 10 µM IVM (blue bar), assessing IVM’s agonistic activity. **b**, Representative traces illustrating the co-application of 10 µM IVM with pH 9 (grey bar) to evaluate modulatory effects on an uninjected oocyte and an oocyte expressing SsCl_EM_. **c,** Quantification of mean peak currents elicited during pH 9, pH 9 + IVM application, and the peak rebound current that occurs after cessation of pH 9 + IVM application. Data is plotted as mean ± SEM (n ≥ 3 cells per condition). Statistical comparisons were performed using multiple t-tests. Significance levels are denoted as: ns, not significant; * = p ≤ 0.05, ** = p ≤ 0.01, *** = p ≤ 0.001 and **** = p ≤ 0.0001.

In this study, we employ a combination of structural and functional methods to characterize the molecular mechanism of the pH-sensitive SsCl channel and its modulation by IVM. Using single-particle cryo-electron microscopy (cryo-EM), we determine structures of the receptor in closed, desensitized, and open conformations bound to IVM. We use computational tools to predict residues that contribute to pH-sensing, which were functionally explored using site-directed mutagenesis and electrophysiology. Our findings reveal that SsCl adopts the classical architectural fold for a pLGIC however, its desensitization mechanisms surprisingly resemble cationic channels. Furthermore, the pH-sensing region which acts as an agonist-like site does not occur in the traditional orthosteric binding pocket, but in a proximal region of the extracellular domain known to bind cations in other channels^26^.

## Results and discussion

### Structure determination

The pH-sensitive chloride channel identified in the mite *Sarcoptes scabiei* (SsCl)^3^, was engineered to facilitate structural studies. The entire intracellular loop was truncated (131 residues) and replaced with a SQPARAA linker from the prokaryote homologue GLIC, which is the most well-characterised proton-sensitive pLGIC^27,28^ (Sup.Fig. 1 and 3). This construct is termed SsCl_EM_. An N-terminal eGFP fusion variant was also employed to screen for expression and solubilization conditions, by using fluorescence-detection size-exclusion chromatography (FSEC)^29^. This identified n-undecyl-β-D-maltoside (UDM) as the optimal detergent for solubilization (Sup.Fig. 4).

**Figure 3.**
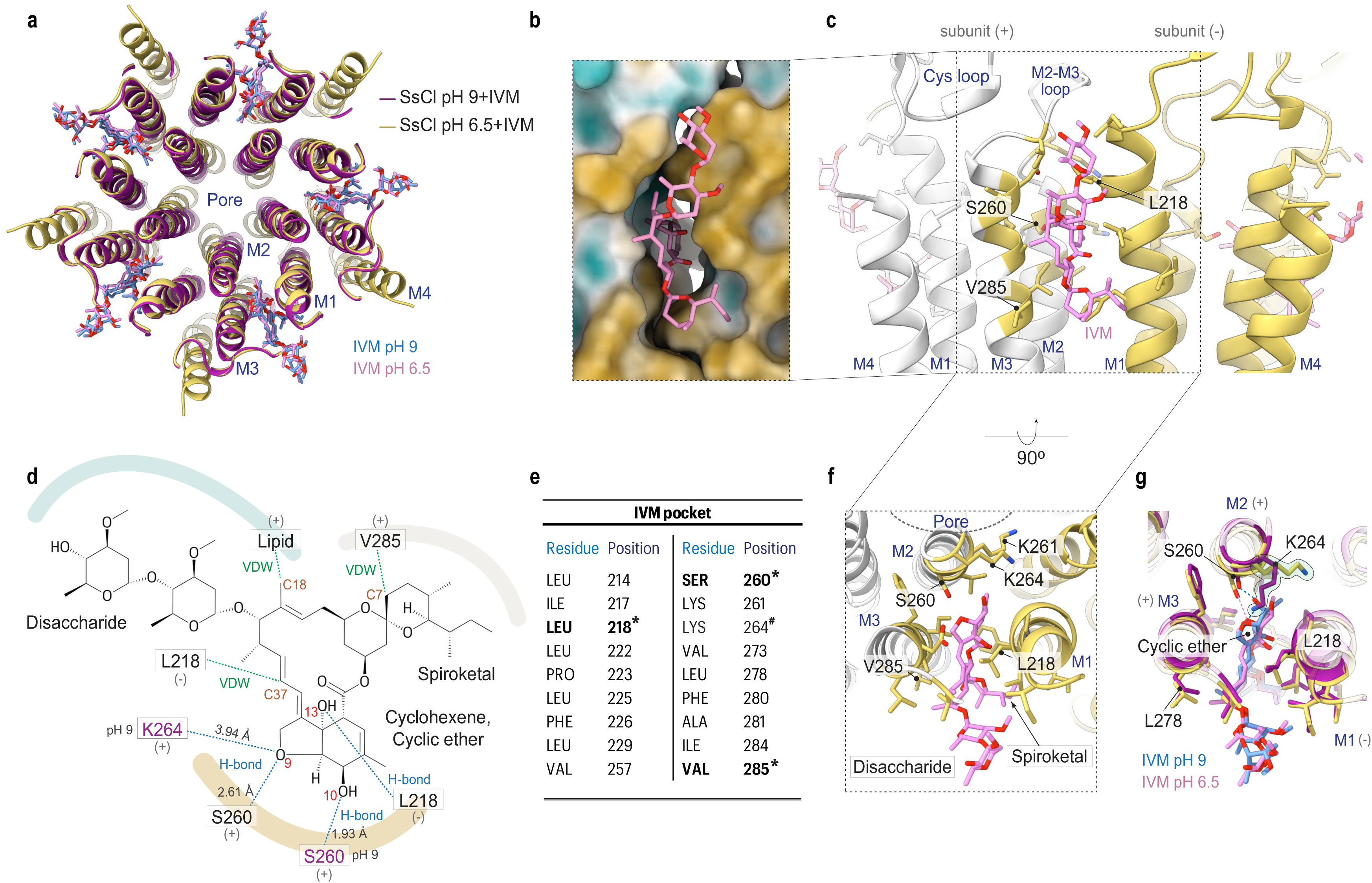
Ivermectin binding site and molecular interactions in SsCl. **a**, Superimposition of SsCl structures at pH 6.5 (yellow and pink) and pH 9.0 (purple and blue) in complex with IVM. **b**, Surface representation of the IVM-binding pocket in the pH 6.5 structure, colored by hydrophobicity (yellow, hydrophobic; white, neutral; blue, hydrophilic) in ChimeraX; IVM is shown in pink. **c**, Side view of the TMD of the structure at pH 6.5 with IVM, showing the residues forming the IVM-binding pocket, and the residues at conserved positions that interacts with IVM (S260, L218 and V285). **d**, Two-dimensional chemical structure of IVM showing key molecular interactions with the SsCl channel. Hydrogen bonds (H-bond) are indicated in blue dotted lines, and van der Waals (VDW) interactions in green. The hydrophobicity profile of the pocket is visualized relative to IVM structure. The more solvent-exposed disaccharide region of IVM is located near the hydrophilic region of the pocket, light blue, while the cyclohexene and cyclic ether moieties are buried within the hydrophobic region of the pocket, dark yellow. **e**, Table listing the amino acid residues forming the IVM-binding pocket in SsCl. Conserved interacting residues are marked in bold with an asterisk (*). Lys264, which adopts different orientations at pH 6.5 and pH 9.0, is marked with a hash symbol (#). **f**, Top view of the pH 6.5 structure showing IVM nestled within the binding pocket. **g**, Top view of the superimposed structures at pH 6.5 and pH 9.0 with IVM, showing the distinct conformations of Lys264 (silhouette).

**Figure 4.**
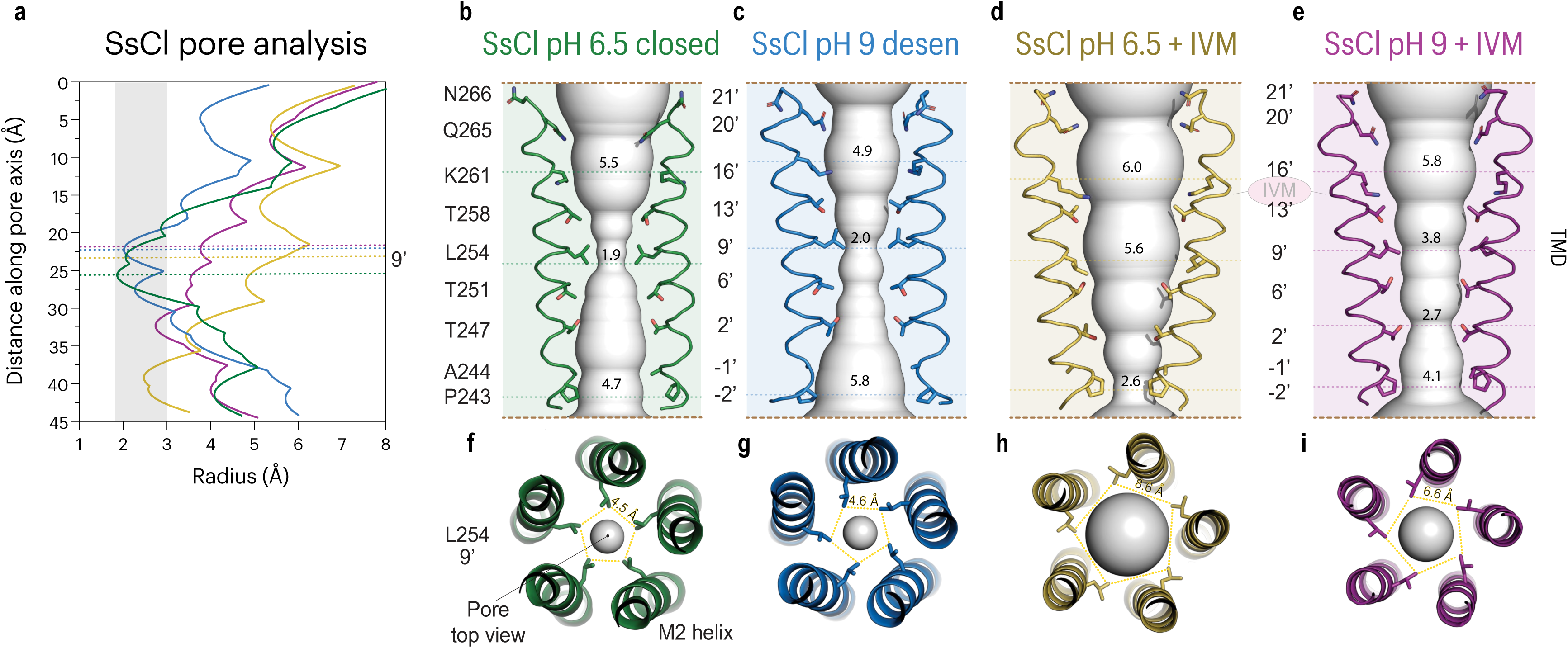
Ion conduction in four SsCl states. **a**, Pore radius profile plotted in Å (x-axis) as a function of longitudinal distance along the channel axis (y-axis), computed using HOLE software. **b–e**, Pore regions rendered as light grey spheres, overlaid with M2 helices and pore-facing residues shown as sticks. Dashed lines mark the 16′, 9′, and –2′ positions, with corresponding radius values indicated. **f–i**, Top views of the pore at the 9′ position, highlighting L254 side chains (sticks). Distances between L254 residues across adjacent subunits are shown with yellow dashed lines.

Functional assays using two-electrode voltage clamp (TEVC) electrophysiology (see methods) confirm that SsCl is activated by alkaline pH. Experiments were also conducted using SsCl_EM_ and produced similar result (Sup.Fig. 5). Therefore, SsCl_EM_ was used as the background construct for all additional experiments. SsCl_EM_ is activated by alkaline pH in a concentration-dependent manner, remaining closed at pH 6.5 with a half-maximal activation (pH_50_) at pH 8.5, similar to previous studies^3^.

**Figure 5.**
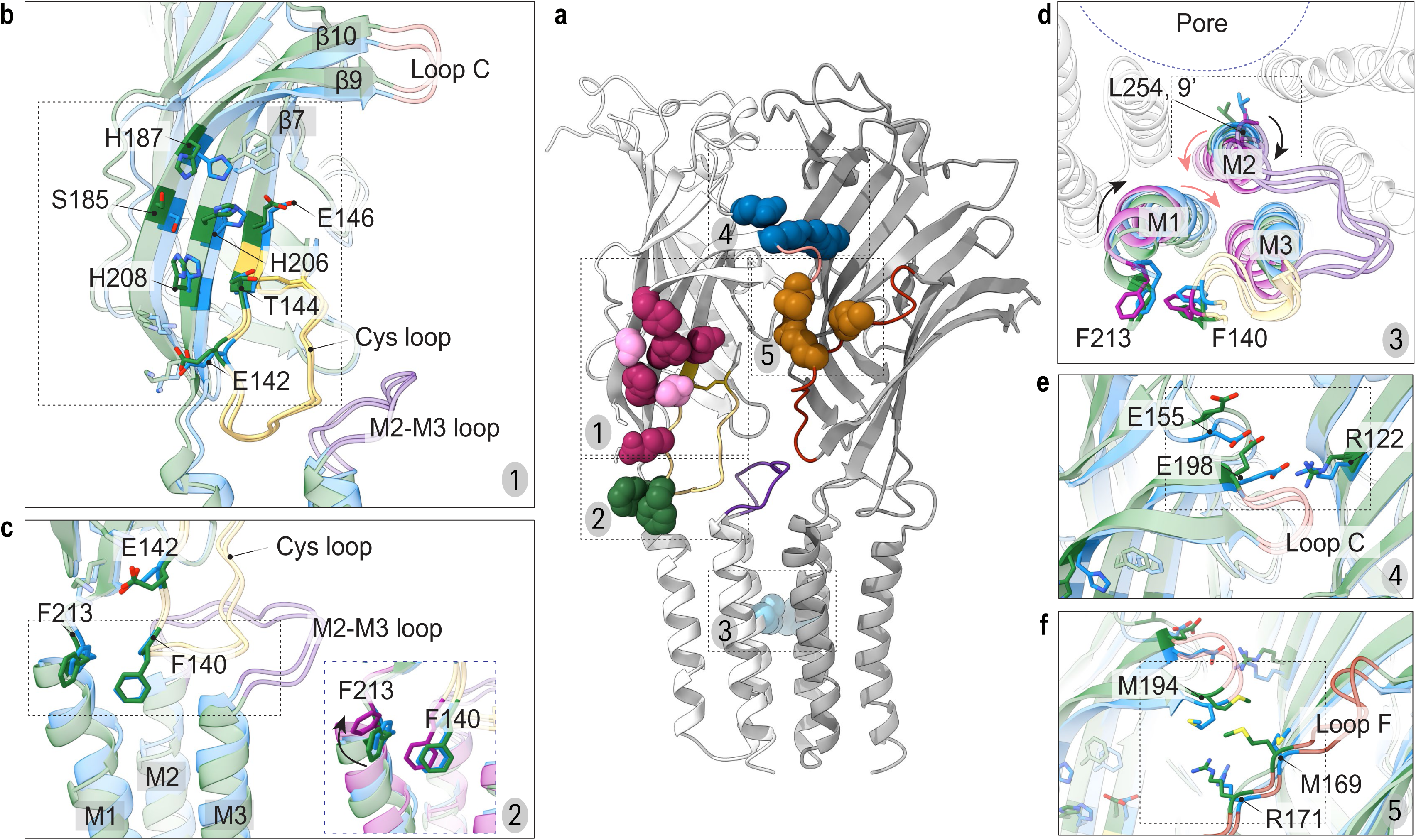
pH-sensing and structural mechanism of channel activation in SsCl. **a**, Side view of the primary (light grey) and complementary (dark grey) subunits, with key residues depicted as spheres: (1) pH-sensor residues (magenta) and additional residues contributing to the pH-sensing network (light pink) in the binding-like region; (2) phenylalanine residues at the coupling interface (green); (3) L254 at the 9’ position of the pore gate (light blue); (4,5) Other residues proximal to the canonical binding region implicated in activation (dark blue and brown). Structural elements are colour-coded: Cys-loop (yellow), M2–M3 loop (dark purple), loop C (salmon), and loop F (dark red). **b**, Superposition of structures at pH 6.5 (green) and pH 9 (blue), highlighting the residues (sticks) that composed the pH-sensor region. **c**, Coupling region side view showing the phenylalanine residues (sticks, near the E142) proposed to mediate channel gating. Inset: superposition of pH 6.5 (green), pH 9 (blue), and pH 9 with IVM (purple) structures, illustrating aromatic interactions that induce displacement of the M1 and M2 helices, most prominent in the IVM-bound open state. **d**, Top-down view of the superposed structures (pH 6.5, green; pH 9, blue; pH 9 with IVM, purple), with L254 (9’), F213, and F140 shown as sticks. Movements of M1 and M2 helices are indicated by arrows (black, upper segment; light red, lower segment). **e,f**, Superposition of pH 6.5 (green) and pH 9 (blue) structures, highlighting residues near loop C that contribute to the activation mechanism.

To capture different functional states of SsCl, pH 6.5 and pH 9 were selected for structural analysis based on its pH-dependent activation profile. In this study, SsCl structures in four distinct states were determined by cryo-EM (Sup.Fig. 6, Sup.Tab. 1): (1) SsCl pH 6.5, (2) SsCl pH 9, and (3) SsCl pH 6.5 IVM and (4) SsCl pH 9 IVM, resolved in the presence of the allosteric agonist, IVM (IVM) (Figure 1). The estimated resolutions for these structures are 4.2 Å, 3.1 Å, 3.1 Å and 3.6 Å, respectively (Sup.Fig. 7). Cryo-EM density maps were used for the manual model building of a single subunit for each structure. Since SsCl assembles as a homopentamer, an initial subunit was built and refined before applying symmetry to reconstruct the complete pentameric structure. Additional model-building and refinement steps were then performed on the assembled pentamer to further improve structural accuracy and consistency. The high-quality density maps enabled the structure building of nearly the entire protein for all structures, with the exception of a few residues at the N-terminus and/or in the M3-M4 loop (Sup.Fig. 8, 9).

**Figure 6.**
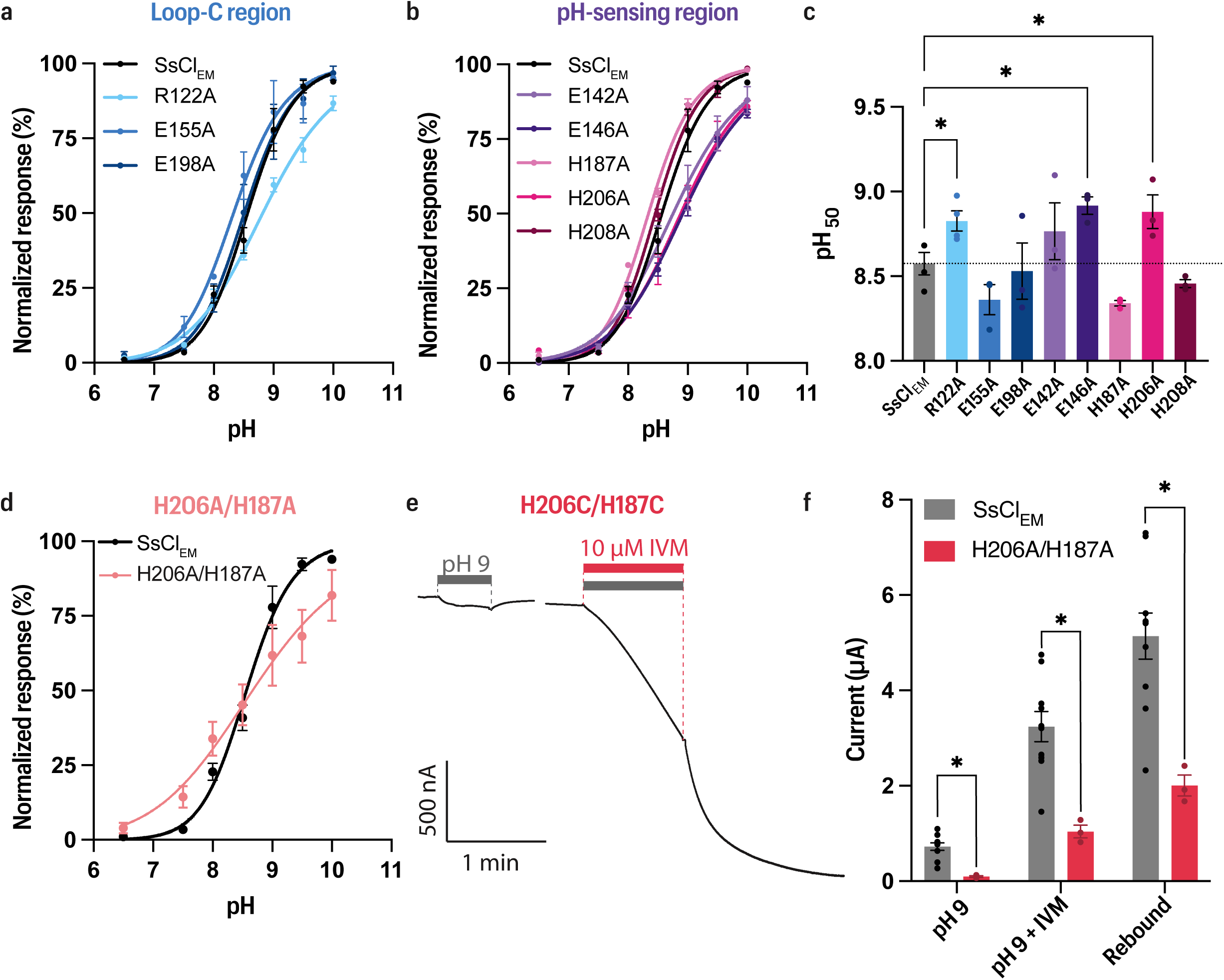
Mutagenesis alters the pH sensitivity of SsCl. **a,** Concentration-response curves for SsCl_EM_ with single-point mutations within the **a,** loop-C region and **b,** pH-sensing region. Data are presented as mean ± SEM (n ≥ 3) and fitted using a modified Hill equation with a variable-slope linear regression model. **c,** Mean pH_50_ values calculated from linear-regression models of individual cells. Data are presented as mean ± SEM (n ≥ 3) and analysed using a one-way ANOVA with Fisher’s LSD test. **d,** Concentration-response curve for H206A/H187A. Data are presented as mean ± SEM (n ≥ 3) and fitted using a modified Hill equation with a variable-slope linear regression model. **e,** Representative traces from oocytes expressing the H206C/H187C, illustrating the effects of pH 9 applied alone or co-applied with 10 µM IVM. **f,** Quantification of mean peak currents elicited during pH 9, pH 9 + IVM application, and the peak rebound current that occurs after cessation of pH 9 + IVM application. Data is plotted as mean ± SEM (n ≥ 3 cells per condition). Values were compared to those obtained for SsCl_EM_ using unpaired t-tests. Significance levels for all panels of the figure are denoted as: * = p ≤ 0.05.

Additionally, in the SsCl structures at pH 9, pH-dependent conformational changes cause the M4 helix to shift closer to the detergent micelle, compromising the resolution in this region. The M4 helix structure could, therefore, not be resolved in the pH 9 structures, however, its presence is visible in the density maps (Sup.Fig. 10). In general, the M4 helices are flexible and highly exposed to the detergent micelle, leading to lower-resolution densities for this region. Despite this, the M4 helices were built for structures at pH 6.5 (Sup.Fig. 8a, b). For all structural analyses, the unsharpened cryo-EM raw density maps were examined to minimize bias.

### Architecture of SsCl channel

As observed for all members of the pLGIC family, SsCl forms a cylindrical-shaped receptor composed of five subunits that form a homopentamer. The symmetrical arrangement of subunits creates a central ion-conducting pore, essential for the channel’s function. Each subunit features a large N-terminus which adopts a twisted β-sandwich structure (β1-β10), constituting the extracellular domain (ECD). A conserved disulfide bond between the β6 and β7 strands, a signature of the Cys-loop receptor family^30^, stabilizes the β-sandwich and aids the overall stability and function of the channel. An additional disulfide bond is also present at the top of the channel, within the first β-sheet. The transmembrane domain (TMD), following the ECD, consists of four α-helical segments (M1–M4) interconnected by loops. The M2 helices line the channel pore and are directly involved in ion conduction and channel’s gating (Figure 1a, Sup.Fig. 11).

In pLGICs, three structural regions coordinate channel activation: (1) the binding pocket within the ECD: including loop C, β-strands of the complementary subunit and loop F; (2) the coupling region at the ECD-TMD interface, including the Cys-loop, pre-M1 segment and M2-M3 linker; and (3) the gating region within the TMD, involving the M2 helices (Sup.Fig. 11). Typically, agonist binding at the orthosteric binding pocket generates structural changes which induces closure of loop C. This initiates a cascade that transmits through the coupling region to the TMD, where rearrangement of the pore-lining helices ultimately leads to channel opening (for review, see ^31^).

SsCl stands apart from conventional receptors as it lacks a classical small-molecule agonist. Instead, its activation is governed by changes in pH. This raises a fundamental question: how do shifts in proton concentration translate into conformational changes that activate the receptor?

SsCl undergoes significant conformational rearrangements in response to different protonation states. Comparisons between the structures obtained at pH 6.5 and pH 9 reveal structural differences within or surrounding loop C, suggesting that this region remains important for the activation mechanisms of the channel (Sup.Fig. 12). Residues in SsCl implicated in pH-sensing have also been identified within the ECD, as described below.

SsCl also features an allosteric binding site for IVM, which closely resembles the IVM-binding site originally identified in the *Caenorhabditis elegans* glutamate-gated chloride channel (GluCl)^22^, indicating a conserved binding mechanism^32^ (Sup.Fig. 13).

### Allosteric activation by IVM and its dependency on pH

IVM is a semi-synthetic compound originating from the discovery of avermectins, a group of natural macrocyclic lactones produced by the bacterium *Streptomyces avermitilis* with potent nematicidal activity ^33^. IVM has been shown to modulate a variety of Cys-loop receptors, acting as a potent allosteric agonist and modulator depending on the specific receptor and context^22,32,34–36^

The SsCl_EM_ construct used for structural analyses (Sup.Fig. 3) was expressed in *Xenopus laevis* oocytes and functionally characterized using TEVC. IVM was shown to activate the channel at pH 6.5 and produce positive modulation it at pH 9, though with notable differences. At pH 6.5 where the channel remains closed, IVM acts as an allosteric agonist and produces an extremely slow activation of the channel, consistent with previous literature^3^. In contrast, co-application of IVM with pH 9 elicits a robust and rapid current that reaches 448% of the current induced by pH 9 alone, thus significantly enhancing channel opening under alkaline condition (Figure 2). There is an additional rebound current that occurs upon the return to baseline pH (pH 5.35) that is greater than the co-application of both agonists. This rebound reaches 711% of the current induced at pH 9 alone (Figure 2). IVM is known to act pseudo-irreversibly at pLGICs and thus, upon the return to pH 5.35, IVM is likely to stay bound for an extended period. This rebound current may be due to IVM transiently stabilising an exaggerated open state that occurs upon the removal of activating pH.

SsCl structures at both pH 6.5 and 9 reveal IVM binding at the subunit interface in the upper TMD. It occupies a pocket formed by M2 and M3 of a principal subunit (+) and M1 of a complementary subunit (-) (Figure 3a). This binding mode closely resembles that observed in the GluCl channel^22^ (Sup.Fig. 13), glycine receptors^35–38^ and α7 nAChR^34^, suggesting a conserved binding mechanism across Cys-loop receptors^22,32^.

The IVM binding pocket is primarily comprised of hydrophobic residues which interact with IVM through van der Waals forces. Two key conserved residues, a hydrophobic L218 and a polar uncharged S260, known for their role in IVM binding in GluCl^22^, are also present in SsCl and contribute to IVM binding and stabilization. L218 contributes with its backbone forming a hydrogen bond with O(13) of the benzofuran moiety of IVM, while its side chain engages in van der Waals interactions with C37, near the benzofuran (Figure 3b-f). In GluCl, a third conserved residue, T285, forms a hydrogen bond with the spiroketal group of IVM^22^. However, SsCl has a valine (V285) at this position, which engages in a van der Waals interaction with C7 of the spiroketal (Figure 3b-f).

Interestingly, a lysine residue (K264) located near the IVM binding pocket alters orientation significantly with pH. At pH 9, K264 orients toward the pocket and likely forms a hydrogen bond with atom O(9) of IVM’s cyclic ether moiety. This interaction draws the IVM’s cyclohexene ring closer to S260, enabling it to form a hydrogen bond with atom O(10) of the cyclohexene. At pH 6.5, K264 adopts an opposing orientation and no longer interacts with IVM. This causes S260 to instead interact with O(9) rather than O(10). These pH-dependent conformational changes result in a stronger interaction between S260 and IVM at pH 9 (1.93 Å) compared to pH 6.5 (2.61 Å) (Figure 3d, e, g, Sup.Fig. 14). This may contribute to the increased efficacy of IVM under alkaline conditions (Figure 2). Moreover, the IVM binding pocket becomes more electronegative under alkaline conditions, specifically in the deepest cavity where the cyclohexene and cyclic ether interact (Sup.Fig. 15). This may also contribute to the pH-dependent effects of IVM.

### Ion-conducting pathway and selectivity

SsCl features a central ion-conducting pathway along its five-fold symmetric axis lined by the M2 helical segments from each subunit. Changes in pH and IVM binding both significantly impact the overall conformation of the pore, leading to distinct ion conduction behaviours (Figure 4, Sup.Fig. 16).

At pH 6.5, the channel adopts an hourglass conformation, with the narrowest constriction occurring in the center of the TMD near position 9’ (L254), with a pore radius of 1.9 Å (Figure 4a, b, f). As a chloride ion has a Pauling radius of 1.8 Å and a radius of 3.2 Å when fully solvated^39^, a pore radius of this size is insufficient for chloride permeation. Accordingly, we assign this structure to the closed state. At pH 9, the narrowest constriction point remains at position 9’ but increases slightly to 2 Å (Figure 4a, c, g), also insufficient for ion permeation. Given that the channel is functionally active at pH 9 (Sup.Fig. 5), yet the structure exhibits a constricted pore, this conformation likely represents a desensitized state resulting from prolonged exposure (>1 day) to activating pH during purification.

When the desensitized structure is viewed from the extracellular side, the M2 helices exhibit a rotational movement relative to the closed state: L254 (9′) rotates clockwise, while P243 (–2′) rotates counterclockwise. This local rotation causes an ‘unwinding’ effect at P243, producing the widest pore radius (5.8 Å) observed among all SsCl conformational states (Sup.Mov. 1). Despite this localized movement the M2 helices largely remain in place, showing minimal translational displacement. Interestingly, the ECD, M1 and M3 helices undergo a concerted compression resulting in a global compaction of the channel that is unique to the desensitized state (Sup.Mov. 2, Sup.Fig. 16).

Strikingly, the desensitization mechanism in SsCl differs from the canonical behaviour reported for other anionic Cys-loop receptors^40,41^. These receptors typically display a V-shaped TMD pore, with a bottom-up constriction-to-expansion profile. In contrast, SsCl at pH 9 adopts an hourglass shape, with its narrowest constriction at the midpoint of the TMD (9’) and expanding toward both the ECD and the bottom of TMD, resembling the closed state (Figure 4b, c). A similar conformation was recently reported for the desensitized state of α7 nAChR, in which the desensitization gate occurs at position 9’ ^34,42^. These findings suggest the desensitized conformation of SsCl closely resembles that of the cationic α7 nAChR, rather than the desensitized conformations reported for other chloride channels.

The impact of IVM on the pore was also examined and was found to significantly increase the overall pore radius at both pHs. This is due to the intercalation of IVM between adjacent M1 (-) and M3 (+) subunits that causes a physical separation of the TMD. Comparing the two structures at pH 6.5, the pore radius at position 9’ increases markedly from 1.9 Å to 5.6 Å in the presence of IVM (Figure 4a, b, d, f, h), which is sufficient to accommodate chloride ions. In the presence of IVM, the channel exhibits its narrowest constriction at the bottom of the pore, specifically at the selectivity filter at P243 (–2’), where the radius narrows to 2.6 Å (Figure 4a, d). Although this is below the radius typically required for a fully solvated chloride ion, we hypothesize that the positive electrostatic potential from the M2 helix dipole may reduce the energy barrier at this site, thereby allowing limited ion permeation. A similar constriction is observed in GlyR structures resolved in complex with glycine and IVM, where the narrowest pore radius also occurs at position –2’. The authors suggest these structures represent intermediate states between open and desensitized conformations^37^. Moreover, the functional data shows that SsCl activation by IVM at pH 6.5 is modest and especially slow (Figure 2). This can likely be attributed to the pore architecture in this state, characterized by a constricted lower region alongside an unusually wide upper pore. Such a configuration may hinder optimal chloride ion stabilization and coordination throughout permeation.

Additional comparisons between the IVM-bound structures reveal that the pore expansion at position 9’ is more moderate at pH 9, where the radius only reaches 3.8 Å, 1.8 Å smaller than at pH 6.5 with IVM. Differences are also observed at position –2′. At pH 9 with IVM, position –2′ exhibits a wider pore radius of 4.1 Å, sufficient for chloride ion permeation. Interestingly, the narrowest constriction shifts upward, occurring at position 2′ (T247) with a radius of 2.7 Å. This shift may contribute to stabilizing and coordinating chloride ions to facilitate ion permeation (Figure 4a, d, e, h, i), which is consistent with functional data showing that IVM more effectively activates the channel at alkaline pH. Taking these observations together, we hypothesized that the mode of IVM action is influenced by pH-dependent conformational changes in SsCl, which in turn affect the architecture and dynamics of the pore. The structural variability aligns with functional data indicating that IVM, in combination with pH 9, results in more efficient channel opening compared to pH 6.5 (Figure 2).

To better understand the structural basis for gating behaviour, the conformational changes at position 9’ were examined across all structures. Position 9’ has long been recognized as a critical determinant in pore gating. In an early 1995 study on the α7 nAChR, the leucines at 9’ were highlighted as key gate-forming residues and named the leucine-ring. The author proposed that bending of the M2 helices toward the central axis of the pore orients the leucines inward, forming a hydrophobic constriction that occludes ion conduction^43^. Recent studies have further reinforced these findings^42^. A similar mechanism is observed in SsCl: At pH 6.5 the L254 side chain faces the channel pore, and upon channel opening with IVM, L254 rotates clockwise (extracellular view) to the side away from the pore axis, relieving the hydrophobic barrier and permitting ion flow (Figure 4 f, h, I, Sup.Fig. 17). Interestingly, in the desensitized state, this position resembles that of the closed state. L254 orients toward the pore, though slightly less pronounced, restoring the hydrophobic gate (Figure 4g, Sup.Fig. 17).

SsCl contains the proline-alanine-arginine motif, a highly conserved feature of the Cys-loop receptor family that plays a key role in the anion selectivity filter^44^. Typically, Pro and Ala residues located at the bottom of the TMD pore are crucial for anion selectivity as described in the literature^22,44^. In SsCl, these residues correspond to P243 (-2’) and A244 (-1’) (Figure 4b-e).

Inspection of the cryo-EM density map of SsCl at pH 9 with IVM revealed a density consistent with a chloride ion within the selectivity filter, near T251 (6′) and above T247 (2′). The distances between the fitted ion and surrounding residues suggest an interaction with the hydroxyl oxygen of T251’s side chain (Sup.Fig. 18). This, together with the relatively constricted pore in this region, may be crucial in coordinating the ion as it moves through the channel. In GluCl, a density corresponding to the chloride ion was also found near to the same position (6’) with similar distances as in SsCl^22^.

To further investigate the selectivity of the SsCl channel for chloride ions, the continuum electrostatic potential was analysed using the APBS (see materials and methods). These calculations were performed for the SsCl structures at pH 9, at the pH in which the structure was determined. Following prolonged exposure to alkaline condition (pH 9), the pore adopts a desensitized state with a constriction at position 9’. This is accompanied by a shift in its electrostatic potential surface where the pore becomes almost entirely electronegative (Sup.Fig. 19a). This creates an unfavourable environment for chloride ion permeation, further supporting the assignment of the pH 9 structure to a desensitized state.

Unexpectedly, for the structure at pH 9 with IVM, the entire ECD and the upper TMD regions exhibit a strong electronegative potential (Sup.Fig. 19b). Among the pore-lining residues within the TMD, only one positively charged residue, K261 (16’), is present (Figure 4b-e). Beyond K261, the electrostatic potential shifts to a more neutral state and becomes increasingly electropositive from T258 (13’) downward. The electropositive potential in the lower TMD arises from the dipole moment of the M2 α-helices that line the pore^45,46^, as the majority of pore-lining residues are polar but not charged (Sup.Fig. 19b).

Taken together, these analyses allow us to define four distinct conformational states of the SsCl channel structures here determined: (1) a closed state at pH 6.5, (2) a desensitized state at pH 9, (3) a partially open state at pH 6.5 in the presence of IVM, and (4) an open state at pH 9 with IVM.

### Structural basis for pH-sensing and channel activation

To investigate the pH-sensing mechanisms of SsCl and identify key residues involved in detecting pH-changes, hereafter referred to as the pH-sensor, the ionizable residues of SsCl at pH 6.5 and 9 were analysed using PROPKA, a tool that predicts p*K*_a_ values of ionizable residues based on 3D protein structures^47^. Residues with lower p*K*_a_ values (≤ 7) and strategically positioned on the structure (*e.g.* accessible to solvent) were selected. From this analysis, five amino acids were predicted to contribute to the pH-sensor (see Methods, Sup.Tab. 2).

The selected residues cluster predominantly within the β7, β9, and β10 strands, forming a binding-like region (pH-sensor). This site is characterized by a central, linear arrangement of three histidine residues H206 (β10), H208 (β10) and H187 (β9), flanked by two glutamic acids E146 (β7) and E142 (β7) and surrounding polar residues which may further support local proton exchange dynamics (Figure 5a,b, Sup.Fig. 20). This region, in particular a conserved histidine analogous to H208 in SsCl, has also been shown to co-ordinate zinc binding in glycine receptors which produces positive modulation of the channel^26,48^.

In SsCl, H206 (β10) and E146 (β7) are predicted to constitute the principal pH-sensor residues and form a key ionic interaction which is sensitive to pH-dependent protonation states. At acidic conditions (pH 6.5), H206 is protonated, allowing it to form an ionic bond with the negatively charged E146. At alkaline pHs (pH 9), H206 becomes deprotonated and electrically neutral, disrupting this electrostatic interaction. This transition likely weakens the structural coupling between β10 and β7, increasing the flexibility of the β7 strand, which contains the conserved Cys-loop disulfide bond. At alkaline pH, H206 and E146 may still interact via van der Waals forces and contribute to the overall structure, however, these interactions are considerably weaker than the original ionic bond (Figure 5b).

Beyond this central interaction, other residues within the pH-sensor, including H208 (β10), H187 (β9), and E142 (β7), are also affected by pH changes through deprotonation and/or altered hydrogen bonding and may play a supportive role. Hydroxyl-containing side chains, such as those of S185 (β9) and T144 (β7), may further help to the local hydrogen-bonding network, also influencing the overall dynamics under different pH conditions (Figure 5a,b).

Disruption or weakening of interactions within the pH-sensor, particularly those within β10 and β7, induces a local clockwise twist at the end of β7 that propagates to the ECD–TMD interface (coupling region) (Sup.Mov. 3). This displaces the Cys-loop which impacts interactions between aromatic residues that appear to be important for conformational rearrangements. For instance, a π–π interaction between F140 on the Cys-loop and F213 on the M1 helix transitions from T-shaped in the closed structure, to a more parallel, off-centered stacking geometry, observed in the opened structure^49^ (Figure 5c). This rearrangement increases the overlap between their π-electron clouds, potentially introducing repulse forces, which consequently drives a clockwise rotation (extracellular view) of the upper M1 helix. This motion is coupled to the adjacent M2 helix (gating region), which undergoes a gear-like rotation. The upper segment containing position 9’ rotates clockwise while the lower segment rotates counterclockwise (extracellular view) (Figure 5d, Sup.Mov. 4). In the structures determined with IVM, the entire TMD undergoes an additional rigid-body clockwise rotation (extracellular view) (Sup.Mov. 5). We propose that these coordinated helical and domain-level motions collectively drive pore expansion, ultimately facilitating channel opening.

Interestingly, the β7 twist occurs not only in structures at pH 9, but also at pH 6.5 when IVM is bound (Sup.Mov. 6). This suggests that the β7 twist is necessary for channel opening, even under acidic conditions where the H206–E146 ionic bond is present. IVM may physically enforce the β7 twist, overcoming the bonds stabilizing effect and contributing to the slower, less efficient channel activation. This contrasts to channel activation at pH 9 where the H206–E146 bond is disrupted, and channel activation is more efficient (Figure 2). We therefore hypothesize that pH-sensing and the associated conformational changes in the presence of IVM, exert a modulatory effect on channel gating, tuning its efficiency rather than acting as a strict binary (on/off) agonist-like mechanism. IVM acts as an allosteric agonist, with its efficacy modulated by the pH-dependent conformational landscape of the channel.

Site-directed mutagenesis was performed to assess the functional contribution of these residues to the channel activity. TEVC recordings reveal that the key pH-sensing residues, H206 and E146, exhibit significantly elevated pH_50_ values compared to WT when mutated to alanine. Interestingly, mutation of H187A and H208 which appear to provide a supporting role in the pH-sensor region, show reduced pH_50_ values; however, these did not reach significance (Figure 6, Sup.Fig. 21). To further explore the importance of the histidine residues for pH-sensing, double-mutations at positions H206 and H187 were examined. When mutated to alanine (H206A/H187A), the channel retains pH-sensitivity; however, the concentration-response curve is greatly impacted due to a reduction in hill-slope value. Interestingly, mutating both positions to cysteine abolishes pH-sensitivity but retains significant currents in response to IVM, including the observed rebound current (Figure 6d-f, Sup.Fig. 21). This highlights the importance of these residues and the flexibility of the β-strands for the pH-sensing mechanisms of SsCl.

Whether this series of protonation-dependent rearrangements represent the unique mechanism for pH-sensing and channel activation remains an open and intriguing question. While the structural cascade centers around H206 and E146, these residues may act in concert with other modulatory elements or alternative sensing pathways, inviting further structural and functional exploration. Notably, these residues are not well conserved across related pH-sensitive channels (Sup.Fig. 1). H206 in β10 of SsCl is predominantly occupied by threonine residues in SecCl channels or charged residues in GLIC and insect pHCl channels. Similarly, the E146 in β7 is predominantly composed of threonine or serine residues, with only GLIC containing a histidine in this position which is not thought to contribute to pH-sensing^50^. This may suggest that SsCl adopts a novel mechanism of pH-sensing compared to related pHCl channels, despite structural and functional similarity.

In addition to the pH-sensor, other structural changes were observed, particularly within the ECD. Comparison of all SsCl structures reveal a significant conformational shift involving β10 (near loop C) of the primary subunit and β6 of the complementary subunit. At the intersubunit interface, the residues E198 (β10) and R122 (β6) form an ionic interaction in the partially opened, opened and desensitized states, which is absent in the closed state. This interaction appears crucial for the movement of loop C and ultimately channel opening. In addition, E155 in the β7–β8 loop bends toward loop C in the partially open, open and desensitized states (Figure 5e, Sup.Fig. 22, 23). Alanine substitution of R122 was shown to significantly increase the pH_50_, supporting its contribution to ECD movements and channel opening (Figure 6). The complementary E155A mutation was conversely found to reduce the pH_50_.

Similar movements in loop C have been reported for multiple Cys-loop receptors and linked to channel activation^31,42^. Loop C typically adopts an outward-facing conformation in the closed state and moves inward to cap the canonical binding site upon agonist binding, or in the case of SsCl, in response to pH changes (Sup.Fig. 24). In SsCl, the ECD also adopts an overall more compact arrangement in the partially open, open, and desensitized states compared to the closed state (Sup.Mov. 7).

Below loop C, an additional hydrophobic interaction is observed between M194 from loop C and M169 from loop F of the complementary subunit. This interaction appears to stabilize the ECD at acidic pH, as it is present only in the structures at pH 6.5. At pH 9, these residues are positioned farther apart, suggesting that the pH-induced conformational changes disrupt this interaction (Figure 5f, Sup.Fig. 22). Furthermore, Loop F exhibits pronounced movement in the partially open and open states, but not in the desensitized state. Specifically, R171 within loop F moves inward, towards the traditional orthosteric binding-pocket (Sup.Mov. 8). Loop F is a critical component of the ligand-binding domain, contributing to the complementary face of the orthosteric site in Cys-loop receptors.

## Conclusion

SsCl is an atypical member of the pLGIC family. It is an anionic chloride-channel activated by alkaline pH in a concentration-dependent manner. SsCl only shares a low sequence conservation with its nearest relatives, such as insect pHCls and the invertebrate GluCl. Despite this, it adopts an architectural fold that is typical for the pLGIC family and shares similar gating mechanism, including a pore constriction at the 9’ position in the closed state. However, a unique feature in SsCl is the location of pH-sensing residues. A collection of non-conserved glutamic acid and histidine residues within the ECD form the pH-sensor and mediate channel activation. IVM can also activate the channel by binding within the TM domains in a similar binding mode to the one observed in GluCl, indicating a common mechanism of ligand recognition. Structurally, IVM greatly widens the pore radius, thus facilitating the flow of chloride ions through the channel. However, distinctive conformations are observed that depend on the environmental pH. At pH 6.5 in the presence of IVM the pore constricts around the selectivity filter, thus limiting ion flow. In contrast, at pH 9 with IVM, the pore is wide open. Another unusual feature in SsCl is the electrostatic surface potential. It is predominantly electronegative in the ECD and upper part of TMD however, becomes neutral about halfway down the pore and is positive near the selectivity filter. The helix dipole effect likely contributes to lowering the energy barrier for ion permeation through the constriction of the selectivity filter. Collectively, these novel findings highlight structural and functional features unique to SsCl, which could be exploited to develop novel and more effective therapeutics targeting scabies.

## Data accession

The atomic coordinates for the four structures - pH 6.5 closed, pH 9 desensitized, pH 6.5 with IVM, and pH 9 with IVM - have been deposited in the Protein Data Bank under accession codes PDB 9RGM, 9RGO, 9RGN, and 9RGP, respectively. The corresponding cryo-EM maps are available in the Electron Microscopy Data Bank under accession codes EMDB-53950, 53952, 53951, and 53953, respectively.

## Experimental procedures

### SsCl cloning, expression and purification

The plasmid encoding SsCl (GenBank ABV02573.1) in the vector pT7TS^3^ was a gift from Joseph A. Dent, MacGill University, Montreal, Canada. For expression screening, we subcloned the SsCl sequence into a modified pFastBac vector containing an N-terminal signal sequence from *Lymnaea stagnalis* AChBP^51^, an 8xHis-tag, GFP, a thrombin cleavage site, a site for ligase-independent cloning (LIC)^52^ and a C-terminal Strep-tag II site. The mature SsCl sequence N28-L489 was cloned into the LIC site and used for initial construct design. To improve protein stability, we truncated the intracellular M3-M4 loop (residues E331-E464) and replaced it with the linker sequence SQPARAA from the prokaryote homologue GLIC^28^. We termed this construct SsCl_EM_. The resulting GFP-fusion construct was employed to optimize expression in *Sf*9 insect cells using the Bac-to-Bac expression system (Invitrogen). Detergent screening was performed using FSEC^29^ and n-undecyl-ς-D-maltoside was identified as a suitable detergent for SsCl solubilization. For large-scale expression, we subcloned SsCl_EM_ in pFastBac as a fusion protein with maltose binding protein (MBP). A thrombin cleavage site was engineered between MBP and SsCl_EM_. Protein expression was induced in 600 mL cell culture volume with an *Sf*9 insect cell density of 2-3 x10^6^/mL. 2 mL of P3 baculovirus was added to the culture and incubated for 4 days at 28°C in a rotary shaker/incubator at 80 RPM.

3.6 L *Sf*9 insect cell culture expressing MBP-SsCl_EM_ was harvested by centrifugation at 10,000 g for 10 min at 4°C. The resulting cell pellet was resuspended in a volume two times the wet pellet weight (v/w) of cell lysis buffer containing 50 mM Na-phosphate buffer and 150 mM NaCl pH 8 supplemented with 1 mM PMSF, 0.1 mg/mL DNase, 0.5 mM MgCl_2_, 1 μg/mL pepstatin, 1 μg/mL leupeptin and 1 μg/mL aprotinin. Cells were lysed by 2 passages through an Emulsiflex C-5 high pressure homogenizer (Avestin) at a maximum pressure of 1500 barr. Membranes were isolated by ultracentrifugation at 125,000 g for 1 hour at 4°C. The membrane pellet was resuspended in 40 mL buffer containing 50 mM Na-phosphate buffer and 150 mM NaCl pH 8. The membranes were solubilized by adding 2% n-undecyl-ς-D-maltoside (Anagrade, Anatrace) for 2 hours at 4°C. The solution was cleared by centrifugation at 30,000 g for 45 min at 4°C. The clear supernatant was loaded on a 3.6 mL pre-washed amylose resin column (New England Biolabs). The column was washed with 10 column volumes wash buffer containing 50 mM Na-phosphate buffer, 150 mM NaCl pH 8 and 0.15% n-undecyl-ς-D-maltoside. The protein was eluted with 5 column volumes elution buffer containing 50 mM Na-phosphate buffer, 150 mM NaCl pH 8, 0.15% n-undecyl-ς-D-maltoside and 50 mM maltose. The eluted fractions were analysed by SDS-PAGE and pooled. 250 units thrombin (Calbiochem) were added to cleave the MBP-SsCl_EM_ fusion protein and incubated overnight at 4°C. The resulting fraction was concentrated to 0.5 mL on an Amicon concentrator with a molecular weight cut-off of 100 kDa. This fraction was then loaded on a Superose 6 Increase 10/300 GL gel filtration column (Cytiva) pre-equilibrated with a running buffer containing 10 mM bis-trispropane at pH 9 or 6.5, 150 mM NaCl and 0.15% n-undecyl-ς-D-maltoside. The eluted fractions were analysed by SDS-PAGE and pooled. The resulting fraction was concentrated on an Amicon concentrator with a molecular weight cut-off of 100 kDa. This solution was then directly used to prepare EM grids. For determination of the IVM-bound structures, 10 μg/mL IVM was added during all purification steps starting from solubilization.

### Cryo-EM sample preparation

Cryo-EM grids were prepared using QUANTIFOIL® R 1.2/1.3 holey carbon films on 300 mesh copper supports. Prior to sample application, grids were glow-discharged for 60 s at 0.39 mBar and 15 mA, using a PELCO easiGlow™ Glow Discharge System. Subsequently, 3 μL of purified protein sample ranging from 2 to 3 mg/mL, was applied to each grid, which was then blotted for either 5.4 or 5.6 s at 4 °C and 95% relative humidity using a Leica EM GP2 automatic plunge freezer. Grids were immediately vitrified by plunge-freezing into liquid ethane and stored in liquid nitrogen.

### Cryo-EM data acquisition

Cryo-EM datasets for the pH 9, pH 9 with IVM, and pH 6.5 with IVM conditions were collected on a Titan Krios transmission electron microscope (Thermo Fisher Scientific) operating at 300 kV with a Gatan BioQuantum energy filter (operated at 20 eV) and Gatan K3 direct electron detector at the Electron Bio-Imaging Centre (eBIC), Diamond Light Source, Harwell, UK. The dataset corresponding to the pH 6.5 (closed state) condition was acquired on a JEOL CryoARM 300 transmission electron microscope (JEOL Ltd.) operated at 300 kV with an Omega filter and K3 direct electron detector, at the VIB-VUB Facility for Bio Electron Cryogenic Microscopy (BECM), Brussels, Belgium. Specifications for data collection can be found in Sup.Table1.

### Cryo-EM data processing

Datasets were processed according to the workflow outlined in Supplementary Fig. 6. Initial processing was performed in RELION 5.0 (ref.^53^). Raw movies were motion-corrected and dose-weighted using RELION’s implementation of MotionCor2^54^, and contrast transfer function (CTF) parameters were estimated with CTFFIND4^55^. Particle picking was conducted externally using crYOLO^56^, employing the general pre-trained model with dataset-specific adjustments to picking threshold and particle size.

Picked coordinates were imported into RELION for particle extraction with initial downsampling to accelerate processing. Extracted particles underwent multiple rounds of 2D classification to remove poorly aligned classes and particles lacking clear structural features of a pentameric channel. Selected particles were used to generate an initial 3D model, followed by unsupervised 3D classification in C1. Particles from well-resolved classes were then re-extracted at the original pixel size.

Further steps were carried out in cryoSPARC^57^. Another 2D classification step was performed, followed by ab initio reconstruction, non-uniform^58^ and local refinement in C1 symmetry. Manual symmetry analysis of the channel confirmed C5 symmetry, which was applied in final rounds of global and local refinement. Global and local CTF refinement were performed to enhance overall map resolution and alignment fidelity. Maps were visualized using UCSF ChimeraX^59^.

### Model building, refinement and validation

An initial reference model was generated using AlphaFold2 (or AlphaFold3^60^) and used as the starting point for model building. Construction began with a single subunit, which was first rigid-body fitted into the cryo-EM density using the ‘fit’ tool in UCSF ChimeraX. Manual rebuilding was then carried out in COOT^61^ to resolve discrepancies and improve the overall fit. Refinement proceeded iteratively within the Doppio suite (CCP-EM^62^), beginning with TEMPy-REFF^63^ to optimize model-to-map agreement. This was followed by successive refinement cycles using Refmac/Servalcat^64^, interspersed with detailed manual building in COOT. Final optimization of local conformations was achieved using ISOLDE^65^ through interactive molecular dynamics flexible fitting. Once the single-subunit model met refinement and validation criteria, C5 symmetry was applied to reconstruct the full assembly. Comprehensive validation was performed using Doppio’s integrated pipeline to assess stereochemistry, fit-to-density, and sequence fidelity. Additional validation of sequence registration was carried out using FindMySequence and CheckMySequence, enabling correction and confirmation of residue-level assignments within the density. Figures were prepared using PyMOL (The PyMOL Molecular Graphics System, Version 3.0 Schrödinger, LLC.) and UCSF ChimeraX. Electrostatic potential was analyzed using APBS^66^.

### Pore analysis with HOLE

Pore dimensions were analyzed using the HOLE program^67^. The SsCl refined atomic models were used as inputs, and the pore axis was defined manually by selecting points along the central channel for each structure. HOLE was run using default parameters to calculate the pore profile, and the resulting radius plot was used to identify constriction sites and estimate the pore diameter along the channel axis. Pore radius profiles were plotted using GraphPad Prism, and structural figures were generated with PyMOL.

### mRNA synthesis for electrophysiology

For expression in *Xenopus* oocytes, the full-length sequence of SsCl was subcloned into pGEM-HE^68^ using *Eco*RI and *Hind*III restriction sites. SsCl_EM_ was engineered the same way as for the pFastBac expression construct. The plasmid was linearized with *Pst*I and transcribed with the mMESSAGE mMACHINE T7 ULTRA Transcription kit. The typical yield of cRNA was around 1.5 μg/μL. Point mutants of SsCl in pGEM-HE were engineered using the QuikChange site-directed mutagenesis kit (Agilent). Mutations were confirmed by Sanger sequencing (LGC Genomics, Germany).

### Oocyte preparation and injection

Ovaries were harvested from *Xenopus laevis* females following procedures established with the Geneva Canton agreement. Ovaries were isolated, placed in sterile medium and kept in clean conditions at 4°C. Ovaries were maintained in physiological OR2 medium (88 mM NaCl, 2.5 mM KCl, 1.8 mM CaCl_2_.2H_2_O, 1 mM MgCl_2_, 5 mM HEPES at pH 7.8, unless indicated otherwise). Oocytes were isolated using mechanical and enzymatic procedures with Type-I collagenase (Sigma). One or two days following isolation, oocytes were injected with 15 nL of mRNA resuspended in distilled water at 0.1 μg/μL. Injections were conducted using the automated Roboinject device (Multichannel System, Germany). Functional expression of SsCl channels was assessed following at least two days after injection.

### Electrophysiology

All recordings were conducted using the automated TEVC system, HiClamp (Multichannel System, Germany). Electrodes were pulled from borosilicate glass (1.2 mm O.D, 0.8 mm i.d.) using a proprietary puller and filled with 3 M KCl, yielded electrodes displaying a typical resistance of about 500 kOhms. Voltage and currents were digitized at 100 Hz and filtered at 20 Hz. Data acquisition and analysis was done using proprietary software running under Matlab (Mathworks Inc.). Unless otherwise indicated, cells were perfused with OR2 adjusted to a pH of 5.35 and maintained at -80 mV throughout the experiment. As desired in the experimental protocol, cells were moved from the perfusion chamber to the incubation well of a 96-microtiter plate (NUNC, Thermofisher). In the microwells, solution was agitated by small specially designed stirring magnet.

## Acknowledgments

This work was supported by an internal KU Leuven C1-grant C14/23/128 and a senior project grant from FWO-Vlaanderen G087921N to CU. JMKF was supported by a post-doctoral mandate (PDM) from KU Leuven PDMt1/24/028. MN is a recipient of a FWO postdoctoral fellowship 12X2722N. CIG is a recipient of a FWO postdoctoral fellowship 1242724N. The Membrane Protein Laboratory is funded by grant 223727/Z/21/Z from the Wellcome Trust with additional support provided by Diamond Light Source and the Research Complex at Harwell, both Instruct-ERIC centres. We thank Diamond Light Source for access to the Cryo-EM facilities at the UK national electron bio-imaging centre (eBIC), proposal nt33941. We also thank Marcus Fislage and Dirk Reiter from VIB-VUB Facility for Bio Electron Cryogenic Microscopy (BECM), Brussels, Belgium, for the support with the data collection of SsCl at pH 6.5 (closed state). The data collection of SsCl at pH 9 (desensitized state) was supported by the iNEXT-Discovery, project PID 30222.

## Author Contributions

J.K.F. and C.U. conceived the project and acquired funding. J.K.F. performed virus production, protein expression, and FSEC; designed the mutants; prepared cryo-EM grids; processed cryo-EM data; built, refined, and validated the models; analyzed structural data; and conducted HOLE and phylogenetic analyses. M.B. prepared bacmid DNA, purified protein, and produced the mutants and mRNA. J.K.F. and P.J.H. collected cryo-EM data under the supervision of A.Q.; P.J.H. also provided intellectual input on cryo-EM data collection and image interpretation. Y.D. conducted electrophysiology experiments, under the supervision of D.B. C.G. performed electrophysiology assays, microscopy of S*f*9 cells, and analyzed electrophysiological data. M.N. contributed to construct design and carried out detergent screening by FSEC. J.K.F. prepared all main structural figures and supplementary material. C.G. prepared electrophysiological data figures and the pLGIC sequence alignment. J.K.F. wrote the original manuscript. J.K.F., C.G., and C.U. were primarily responsible for reviewing and editing the manuscript with input from all authors. All authors reviewed and provided feedback on the final version. C.U. supervised the project.

