## Supplementary data for "Structure of a pH-sensitive pentameric ligand-gated ion channel from the *Sarcoptes* scabies mite"

**Supplementary Figure 1 -**

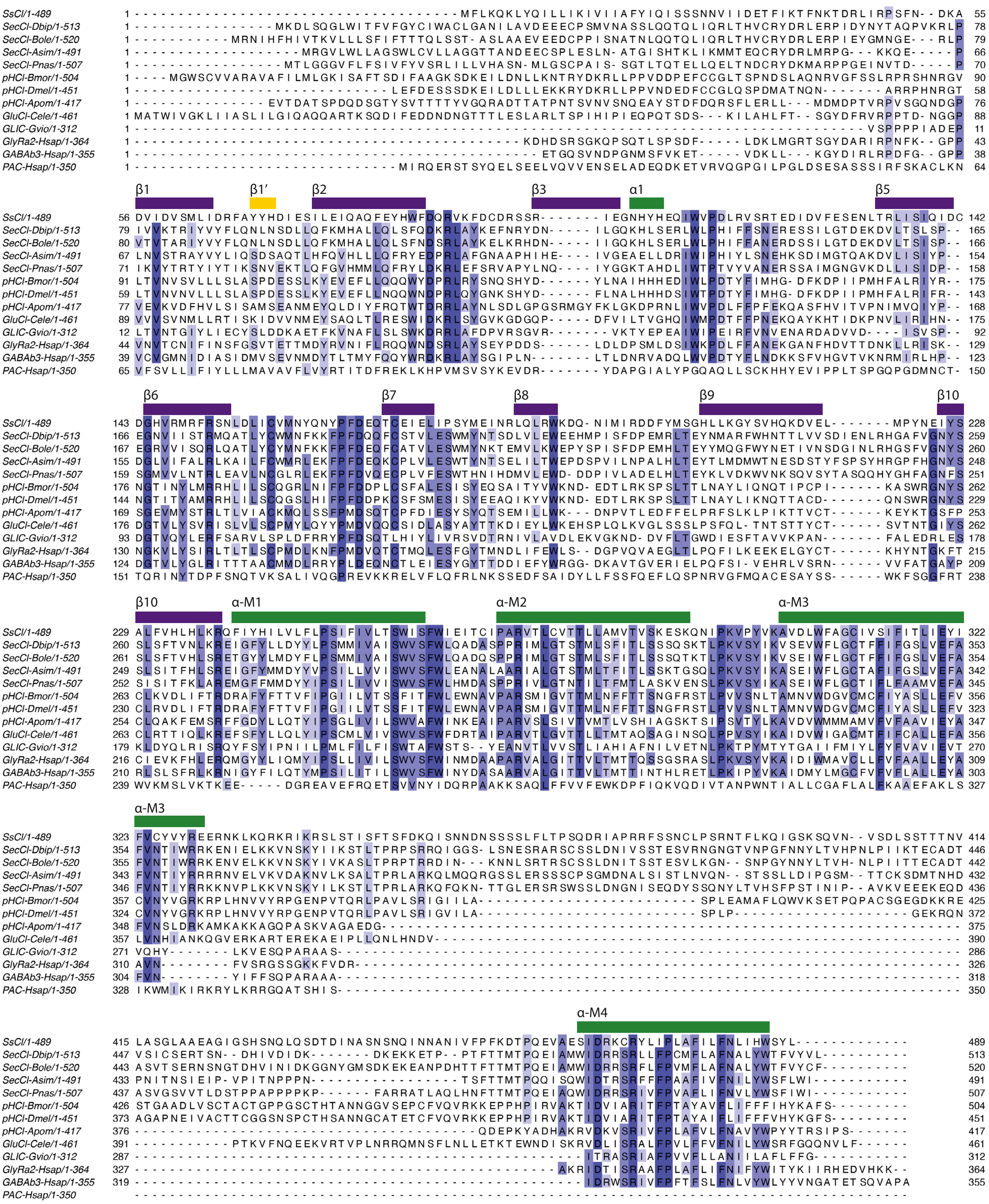

Suplementary Figure 2 -

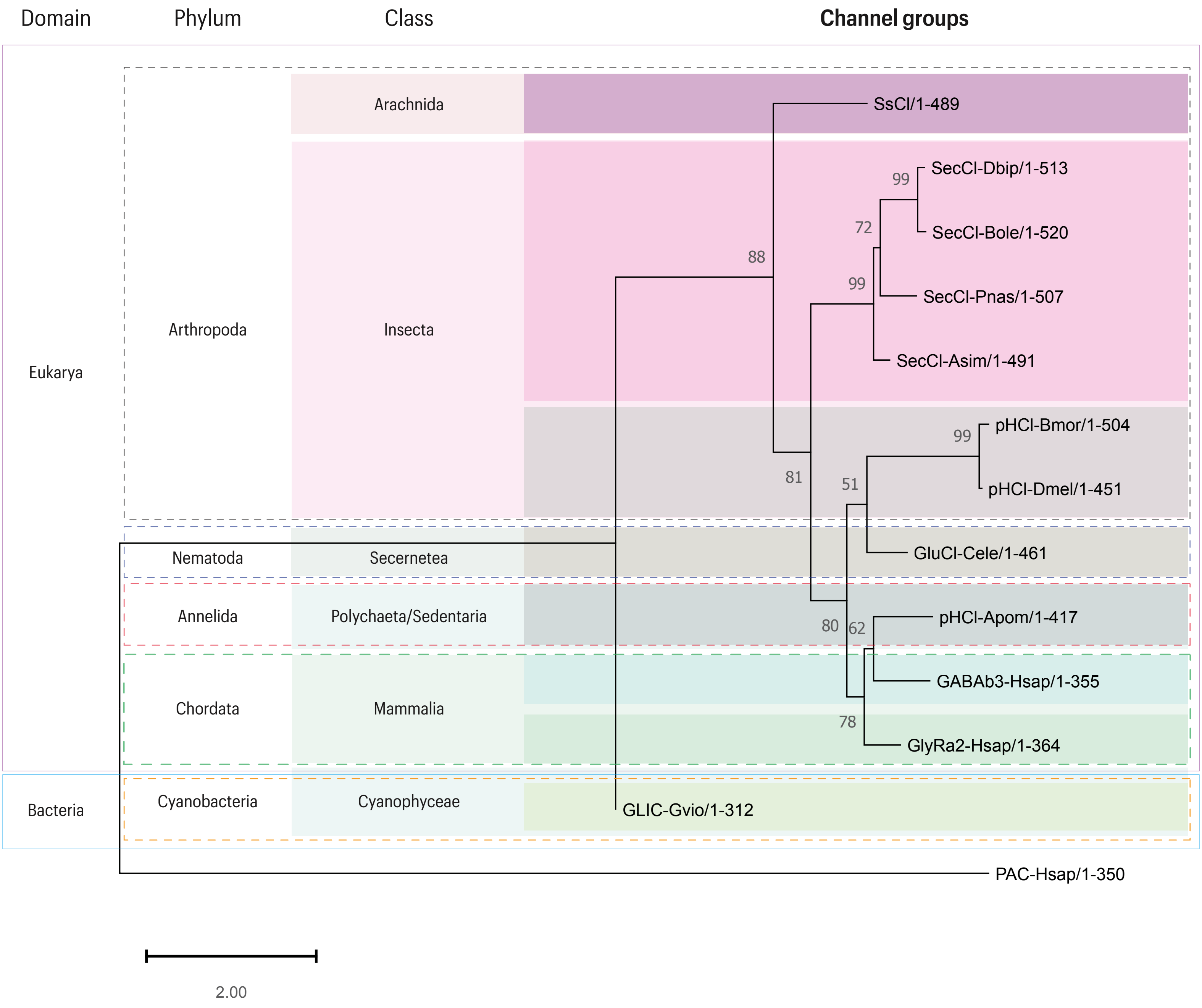

**Supplementary Figure 3 -**

**a**

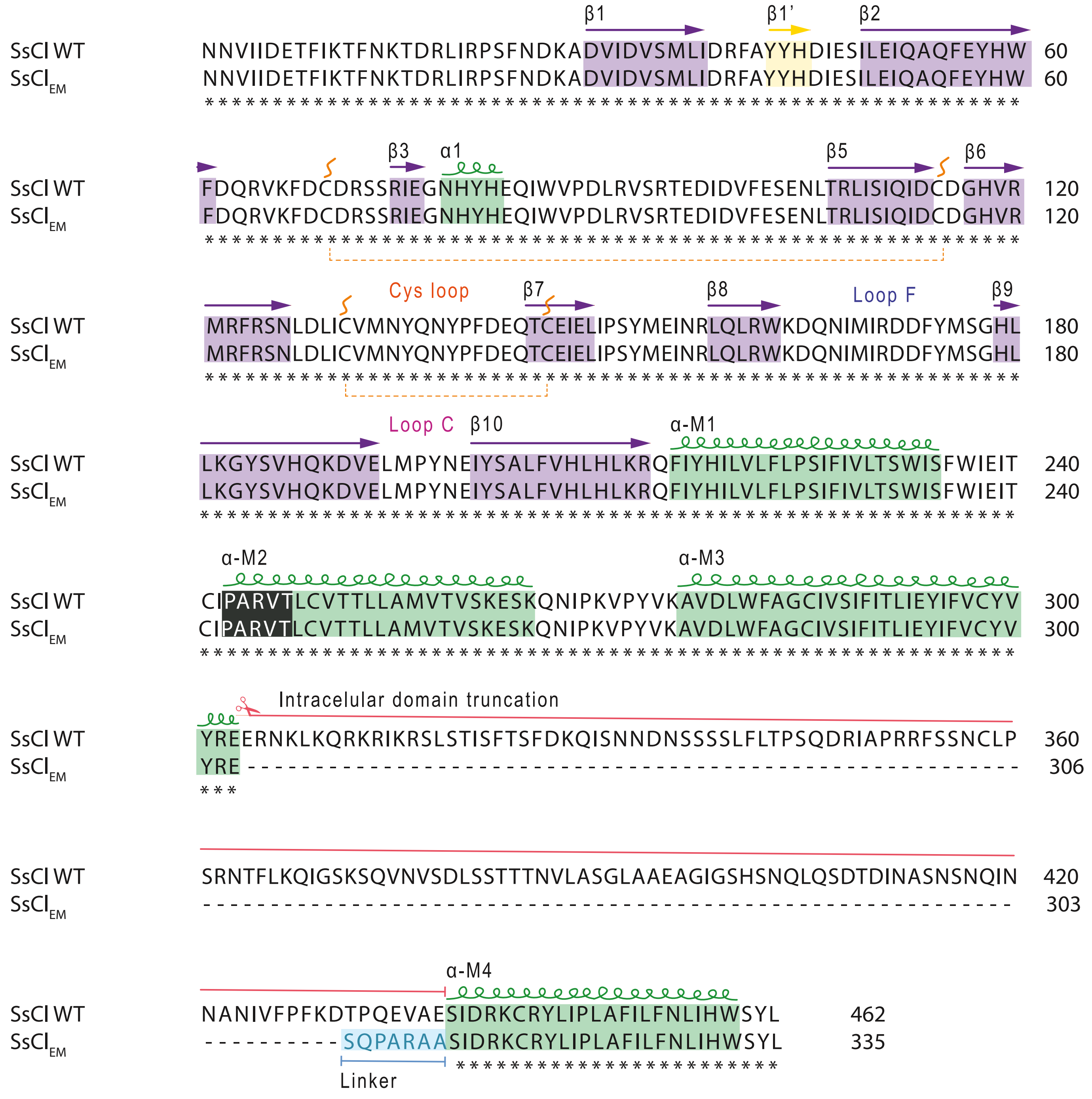


**b**

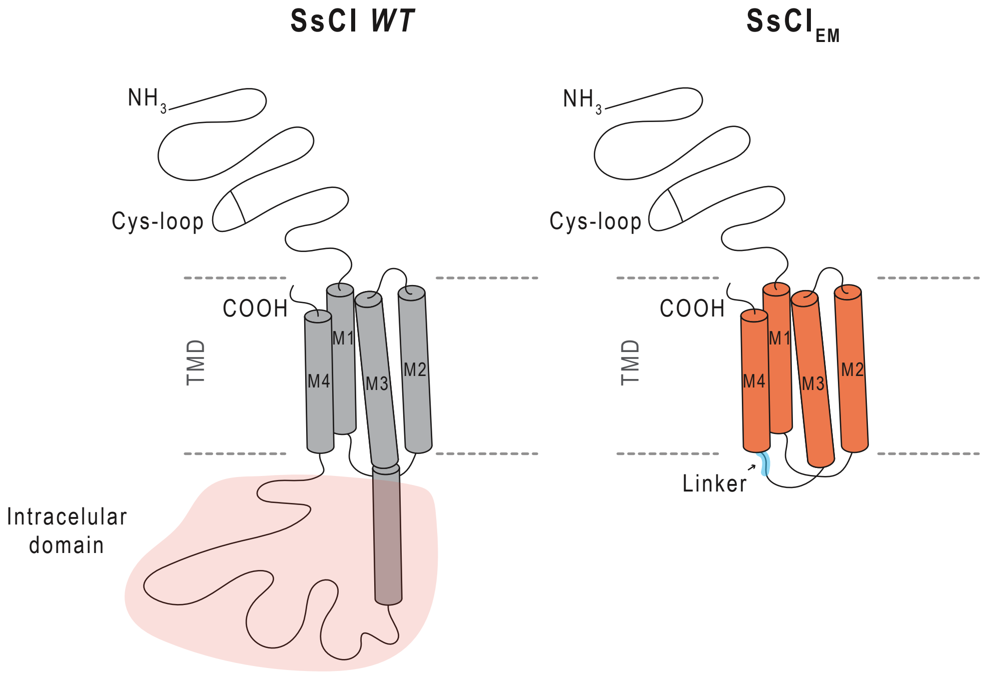

**Suplementary Figure 4 -**

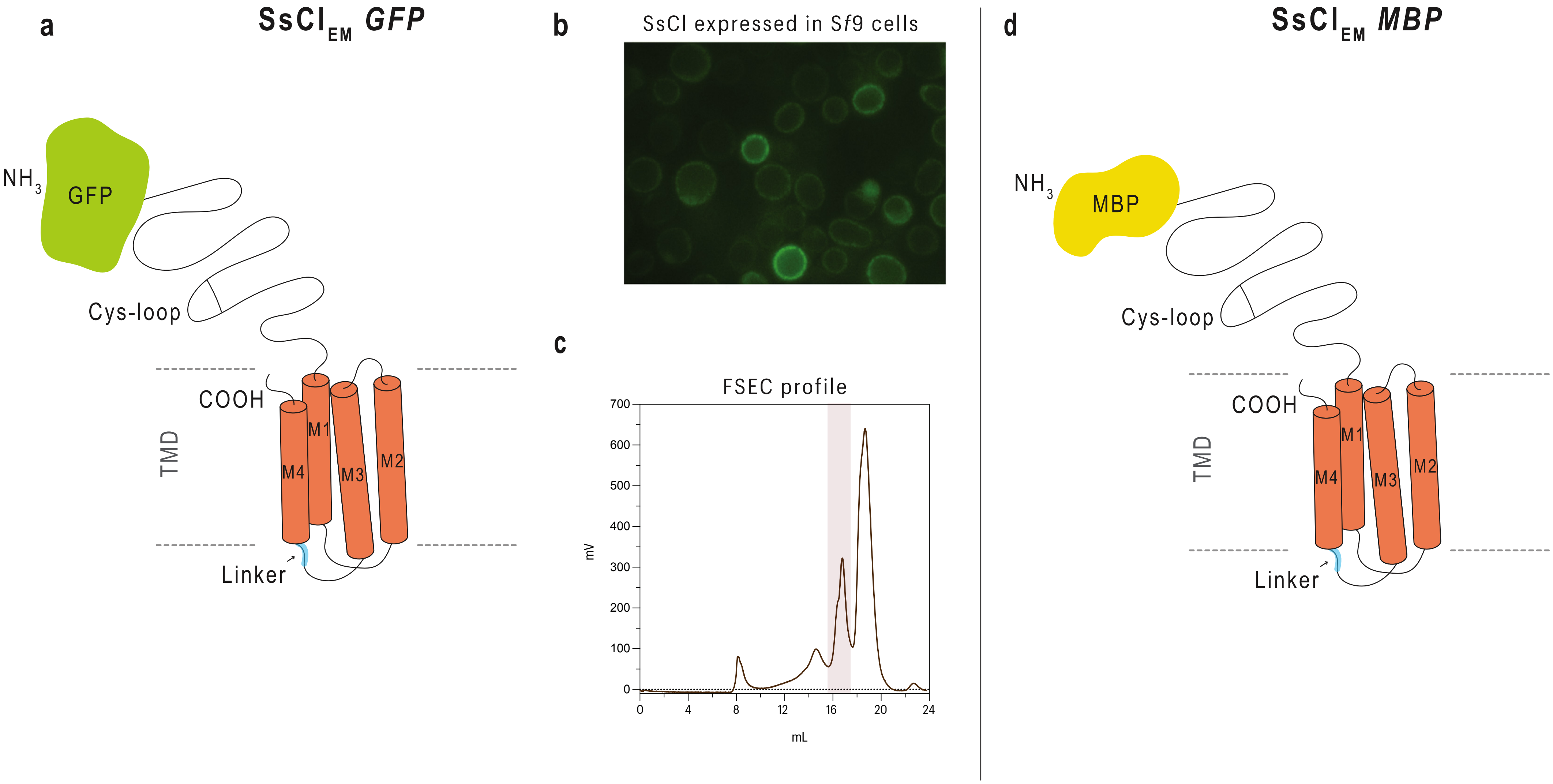

**Supplementary Figure 5 -**

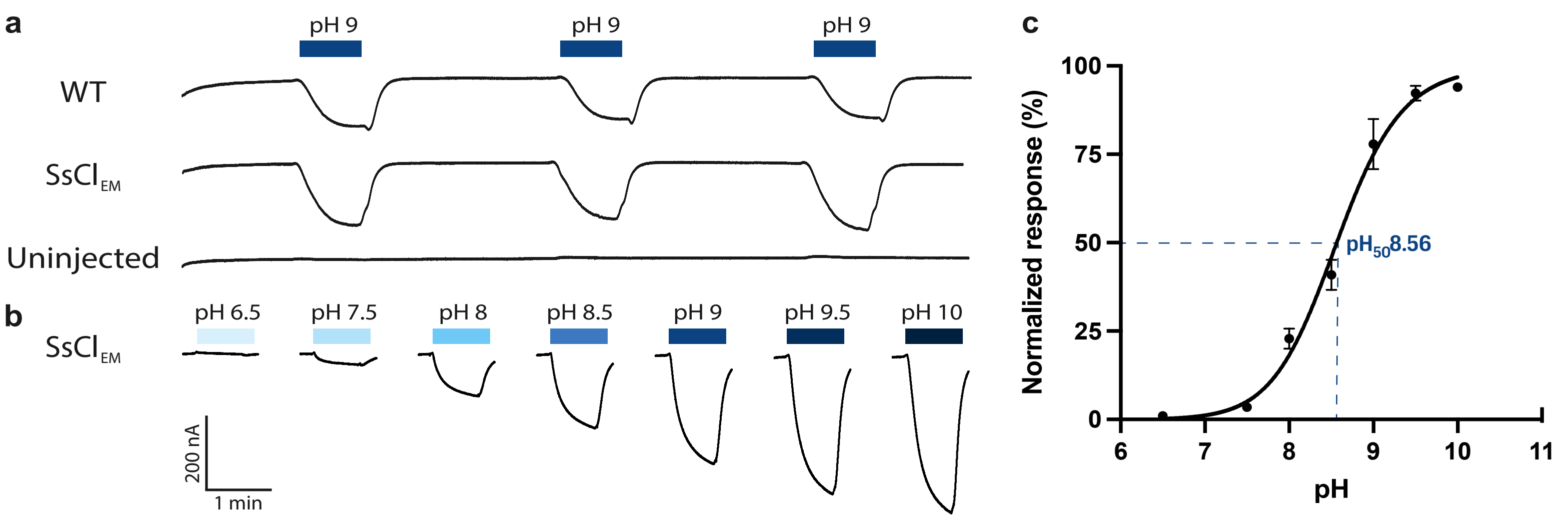

Supplementary Figure 6 -

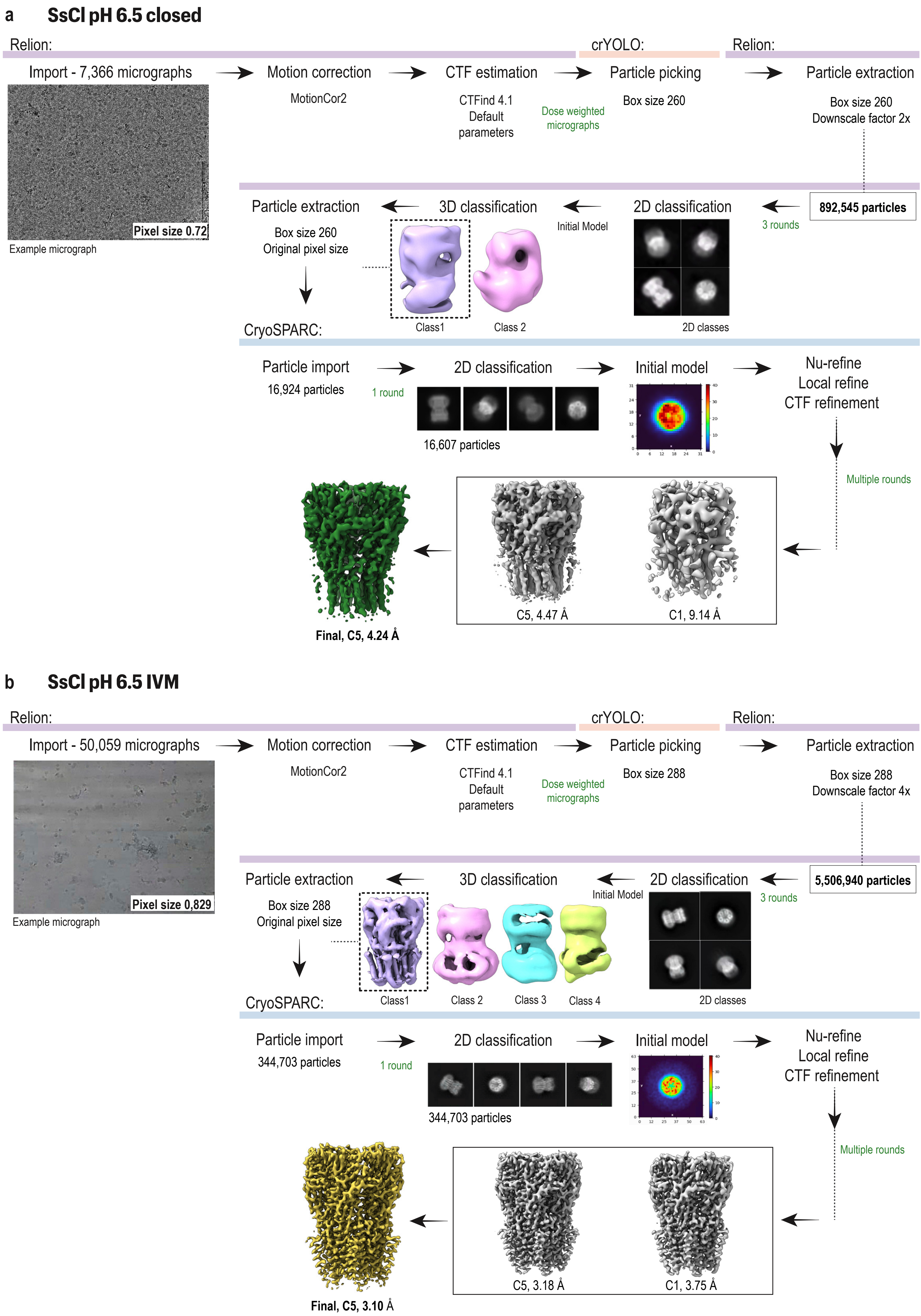

### Supplementary Figure 6 -

#### c SsCl pH 9 desensitized

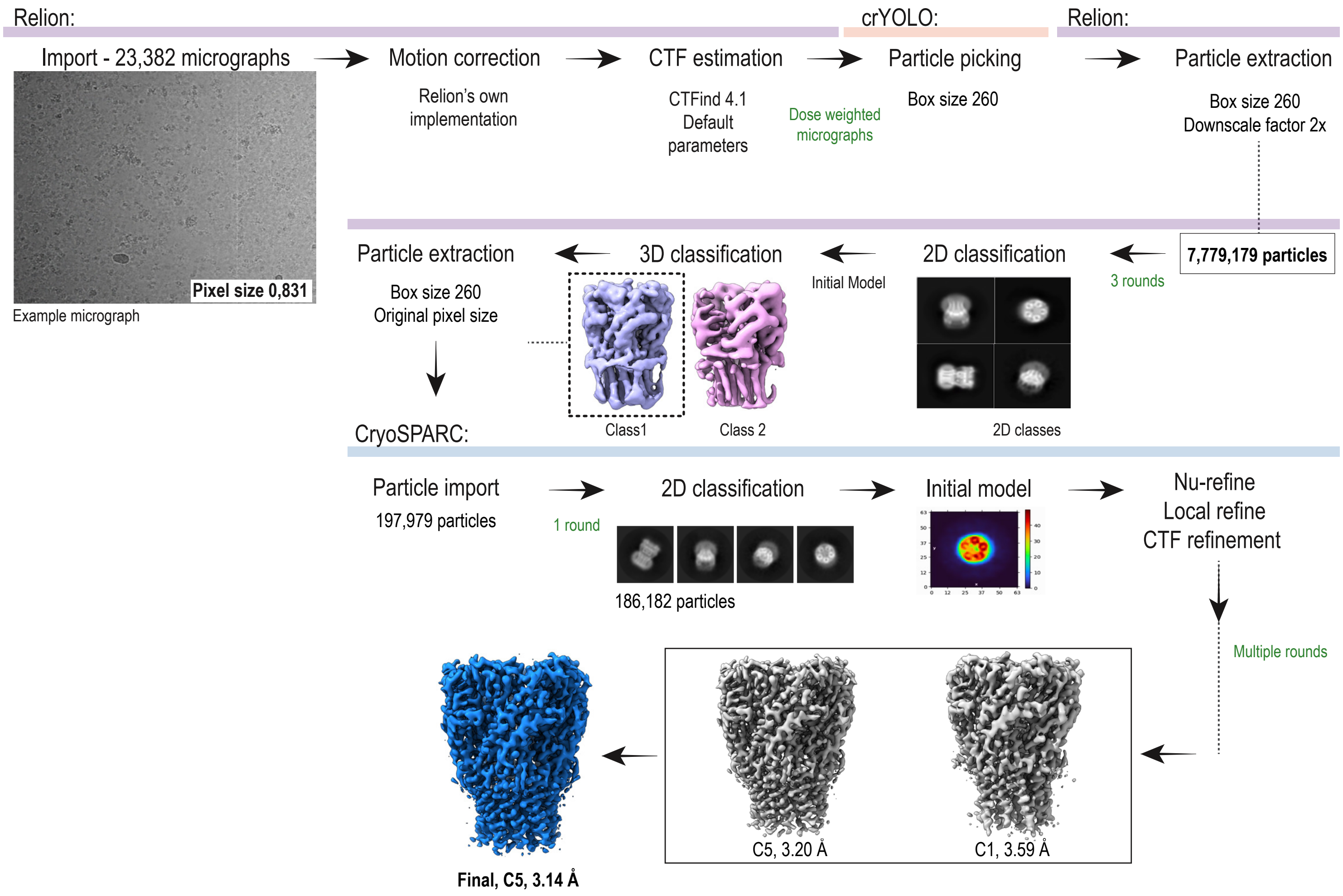


#### d SsCl pH 9 IVM

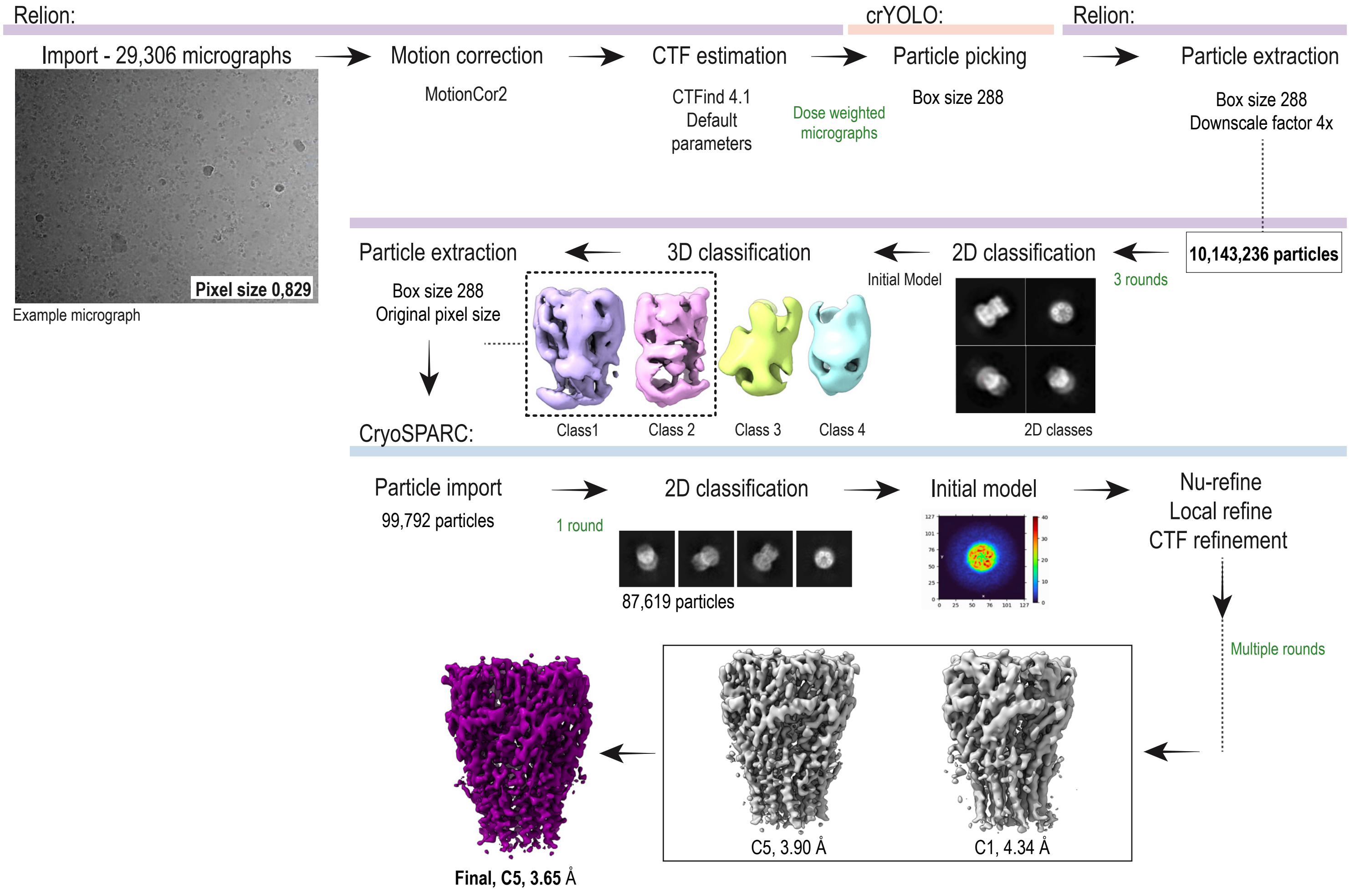

**Supplementary Figure 7 -**

### a SsCl pH 6.5 closed

#### Local resolution estimation

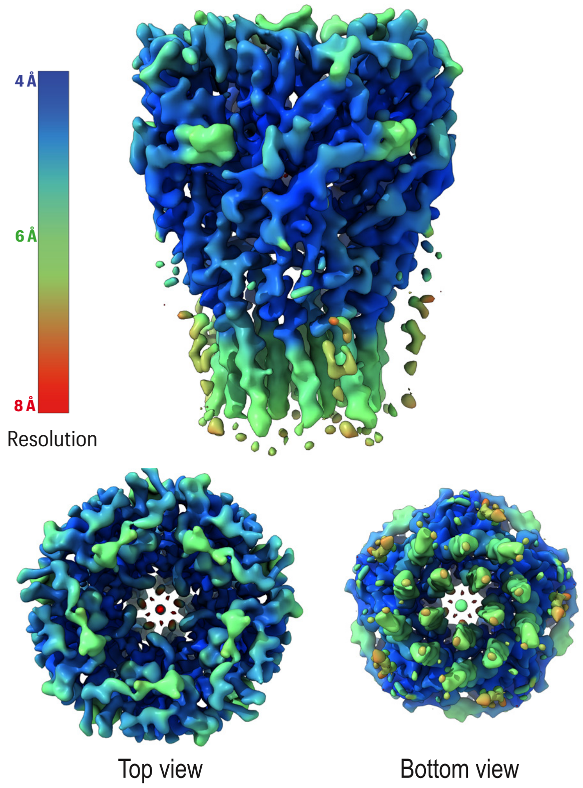


#### Fourier shell correlation (FSC)

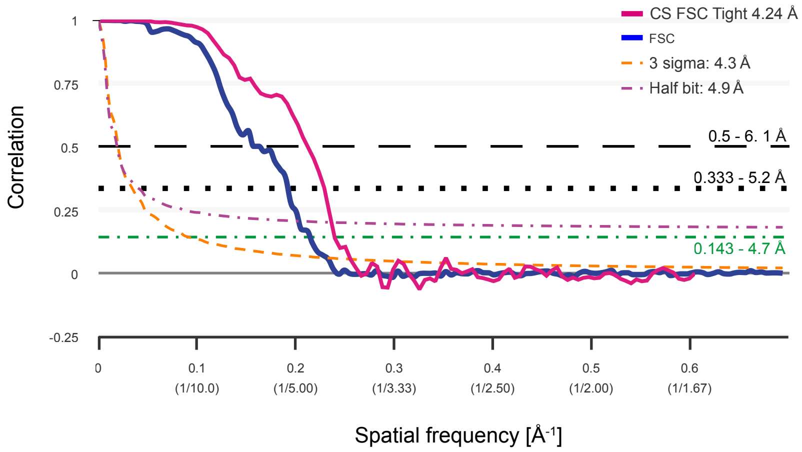


### b SsCl pH 6.5 IVM

#### Local resolution estimation

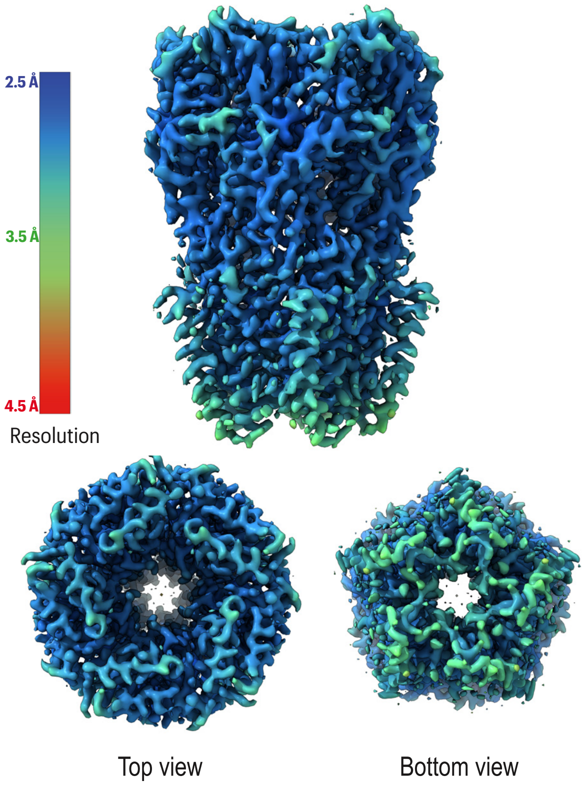


#### Fourier shell correlation (FSC)

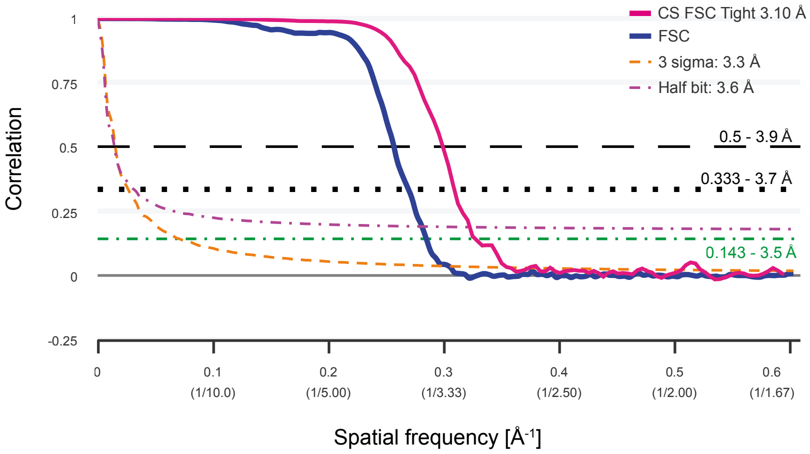

Supplementary Figure 7 -

### c SsCl pH 9 desensitized

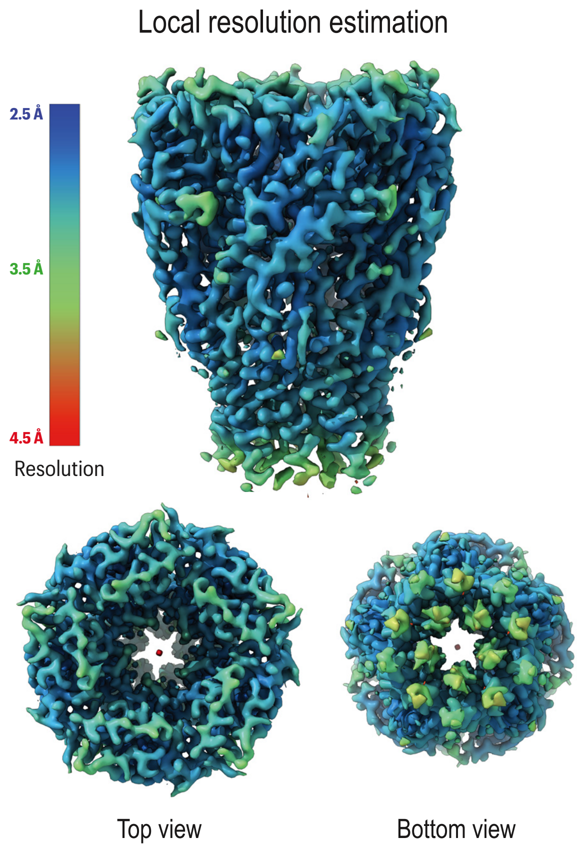


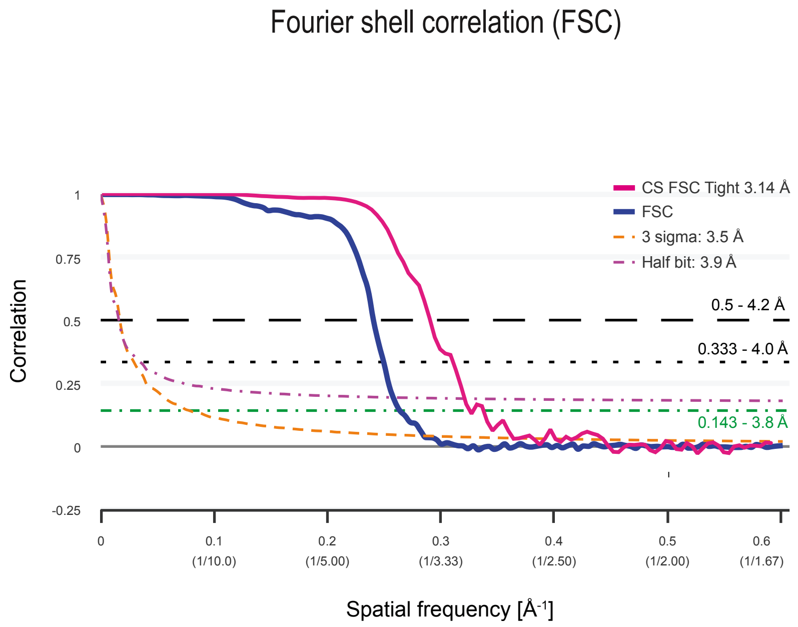


### d SsCl pH 9 IVM

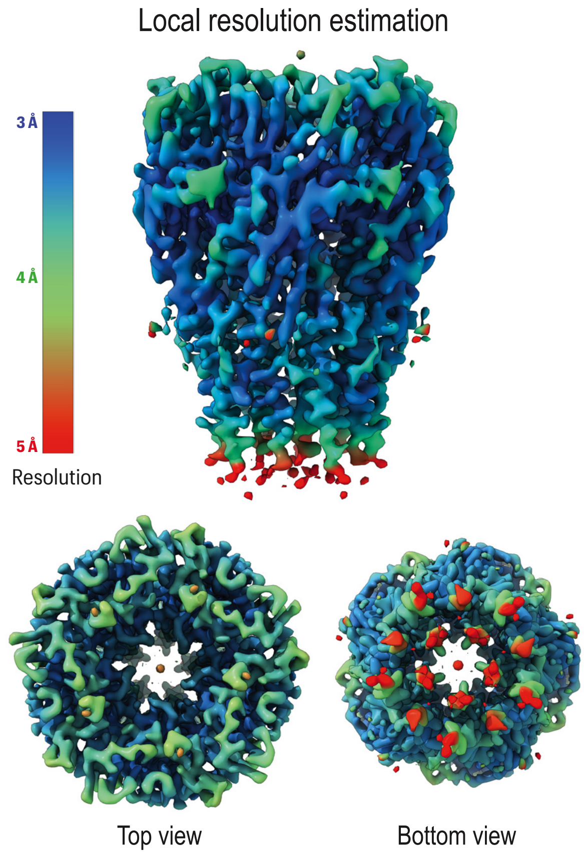


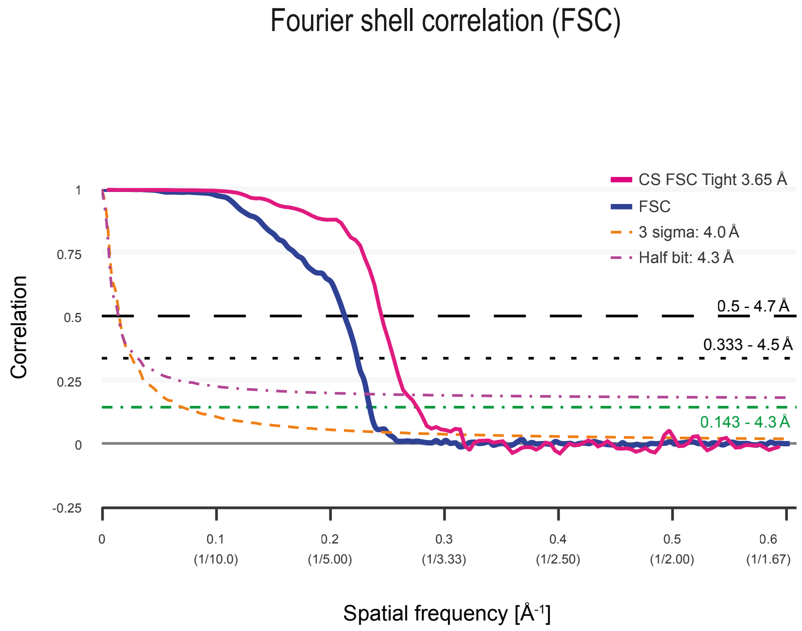

Supplementary Figure 8 -

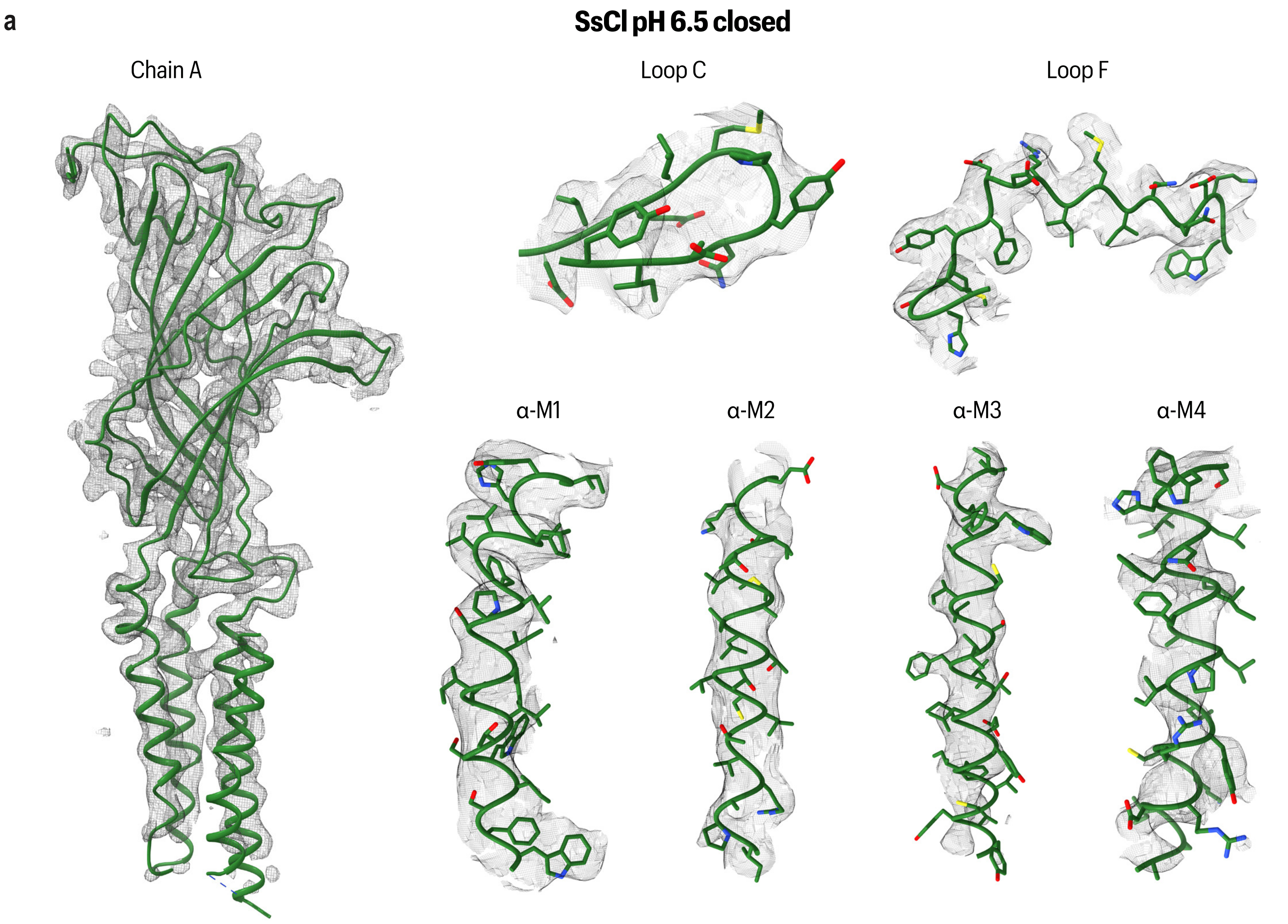


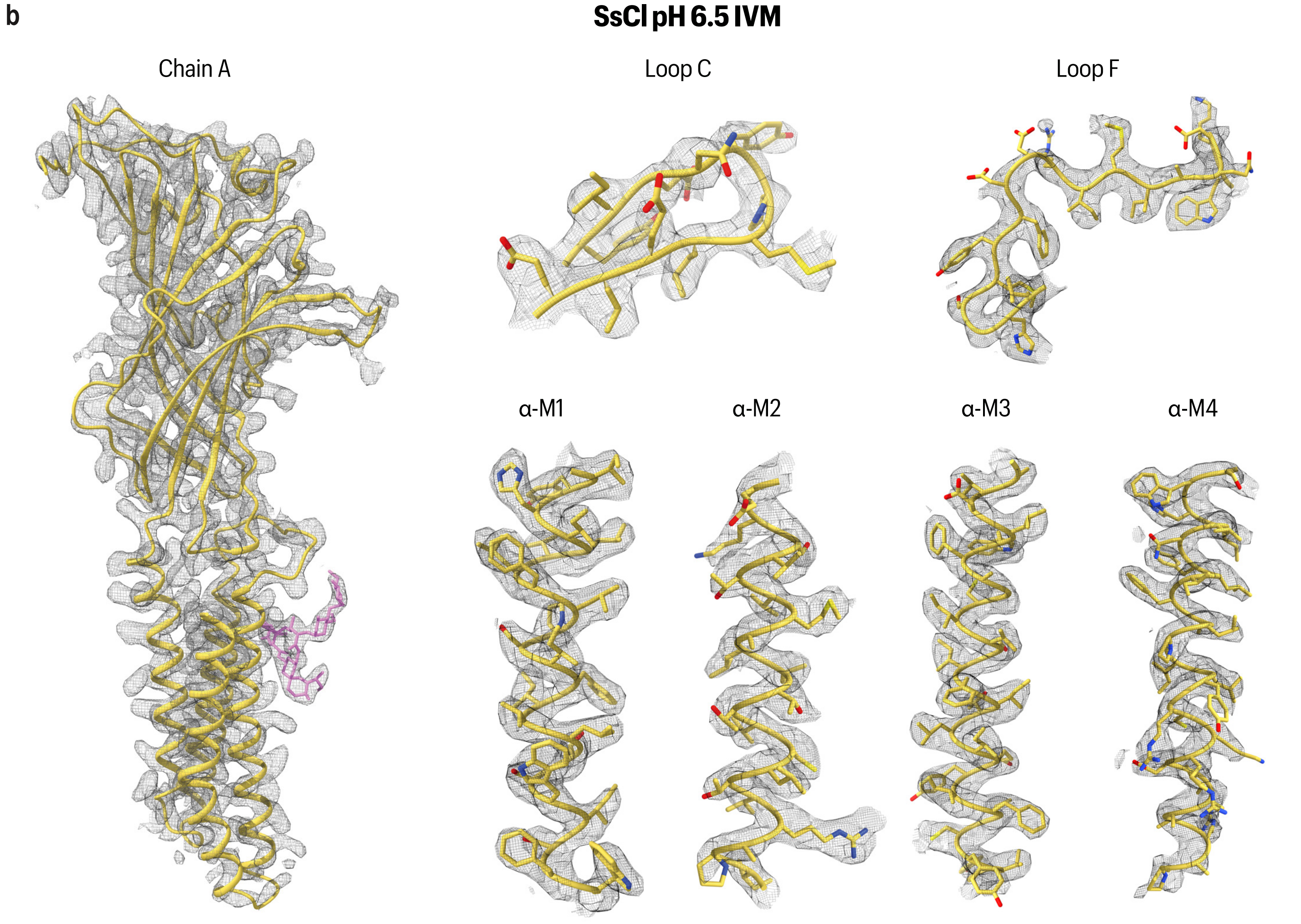

Supplementary Figure 8 -

c

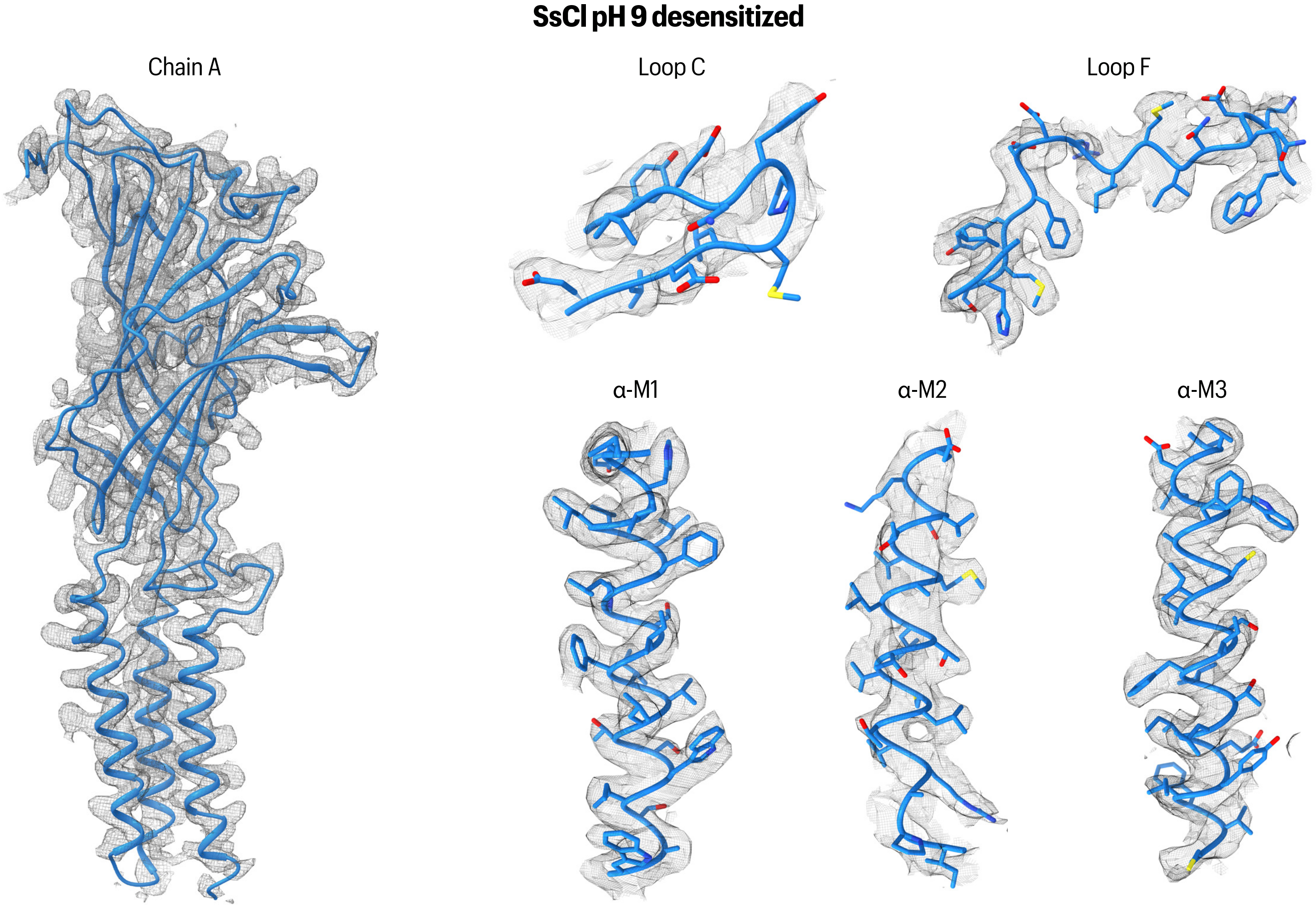


d

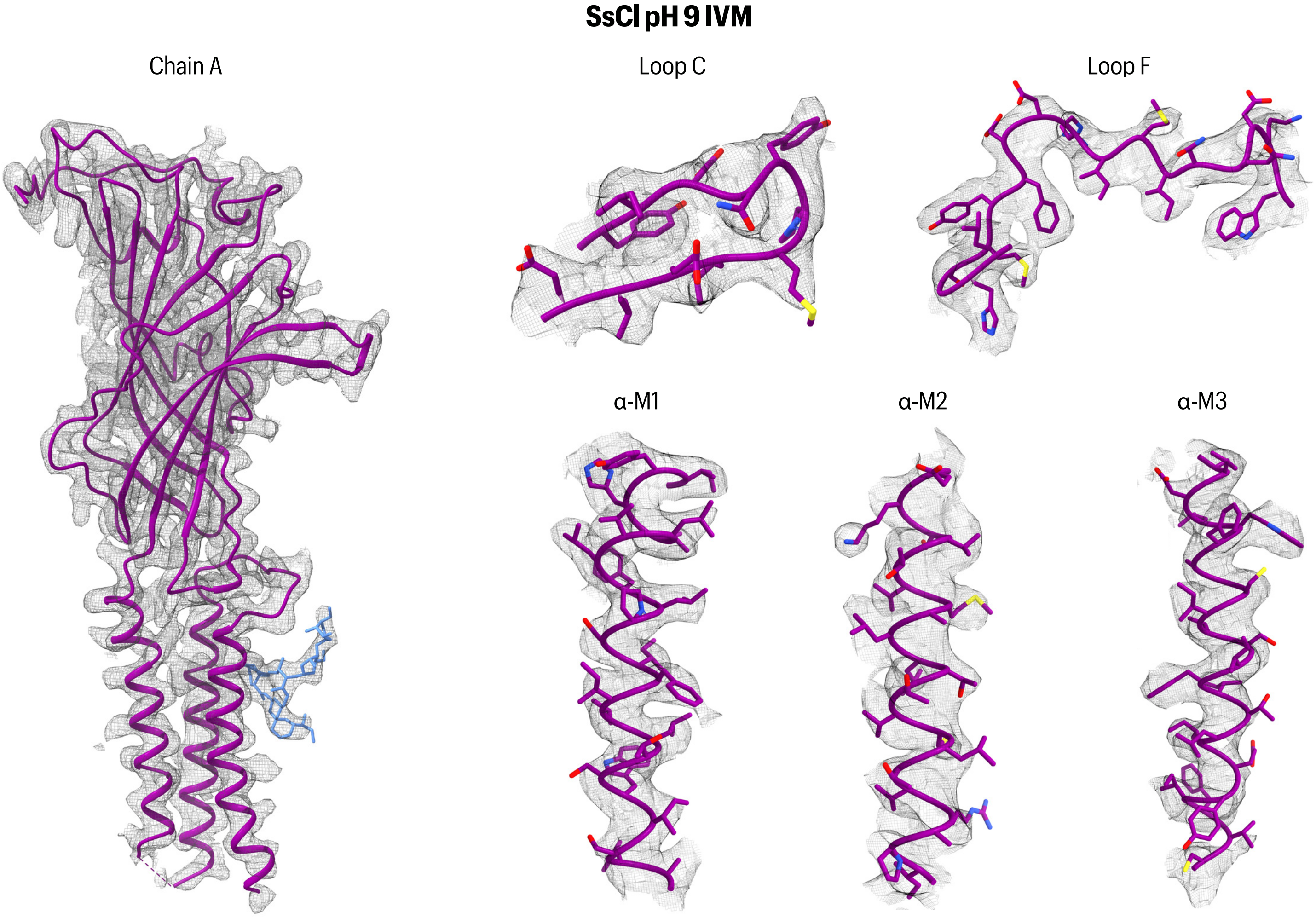

**Supplementary Figure 9 -**

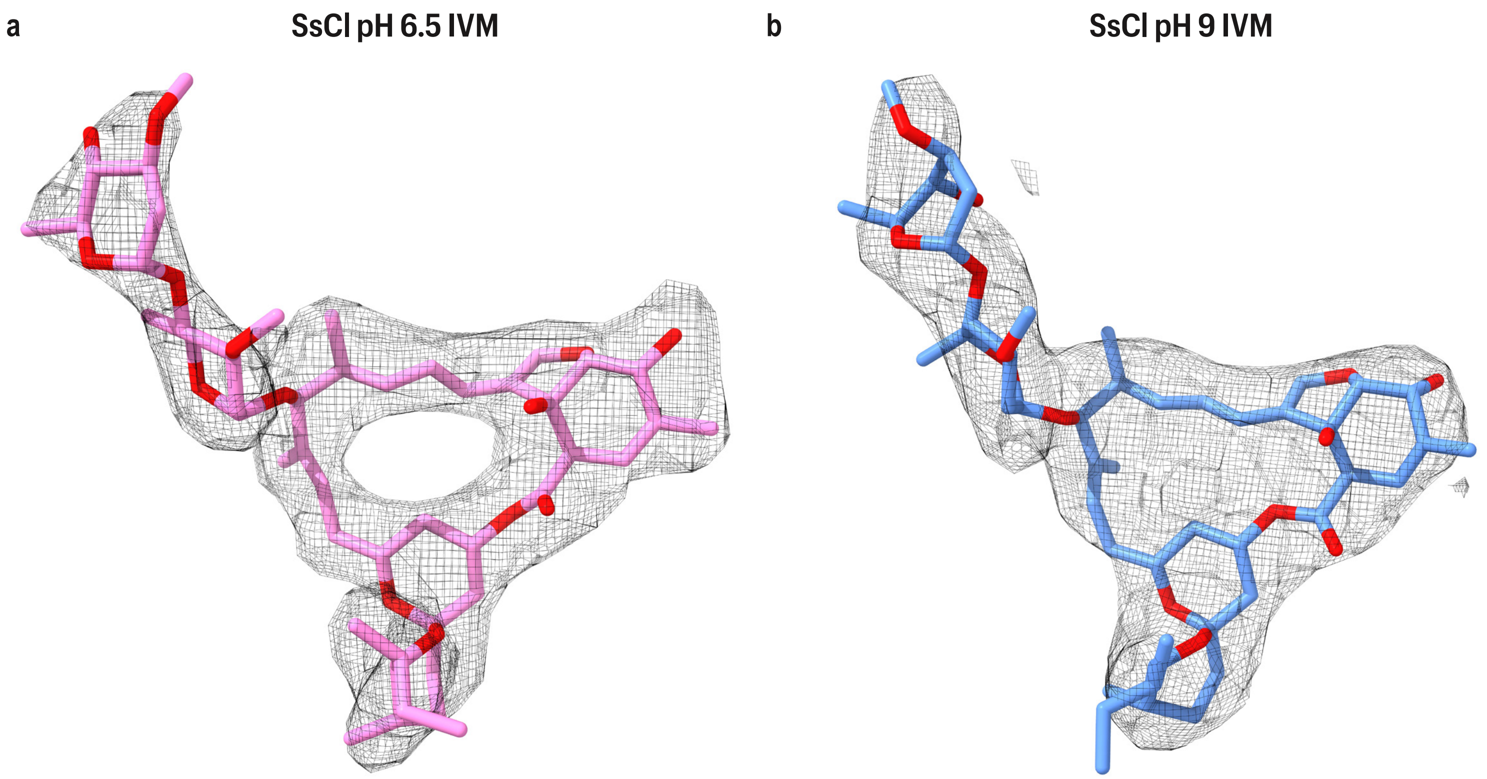

Supplementary Figure 10 -

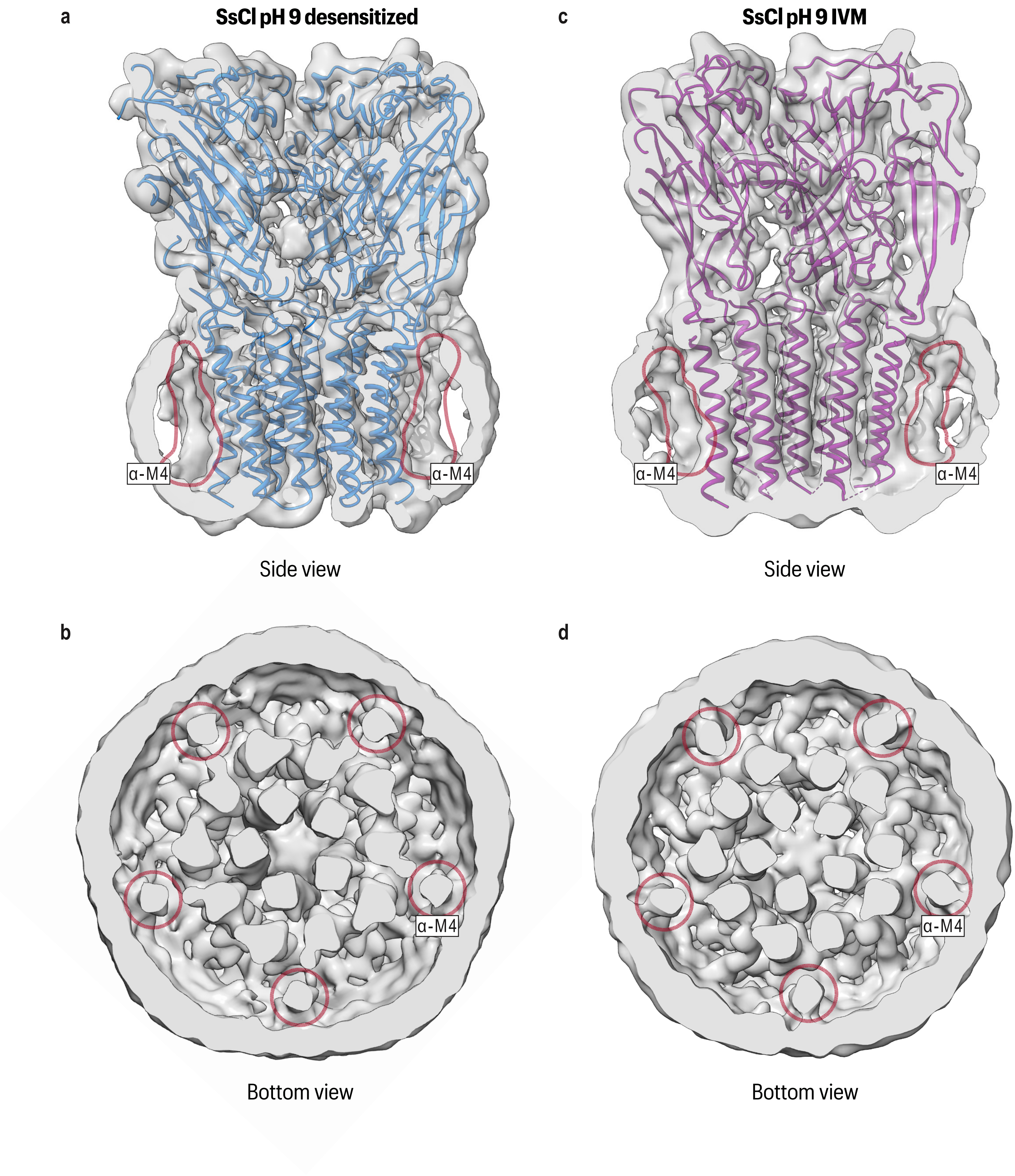

**Supplementary Figure 11 -**

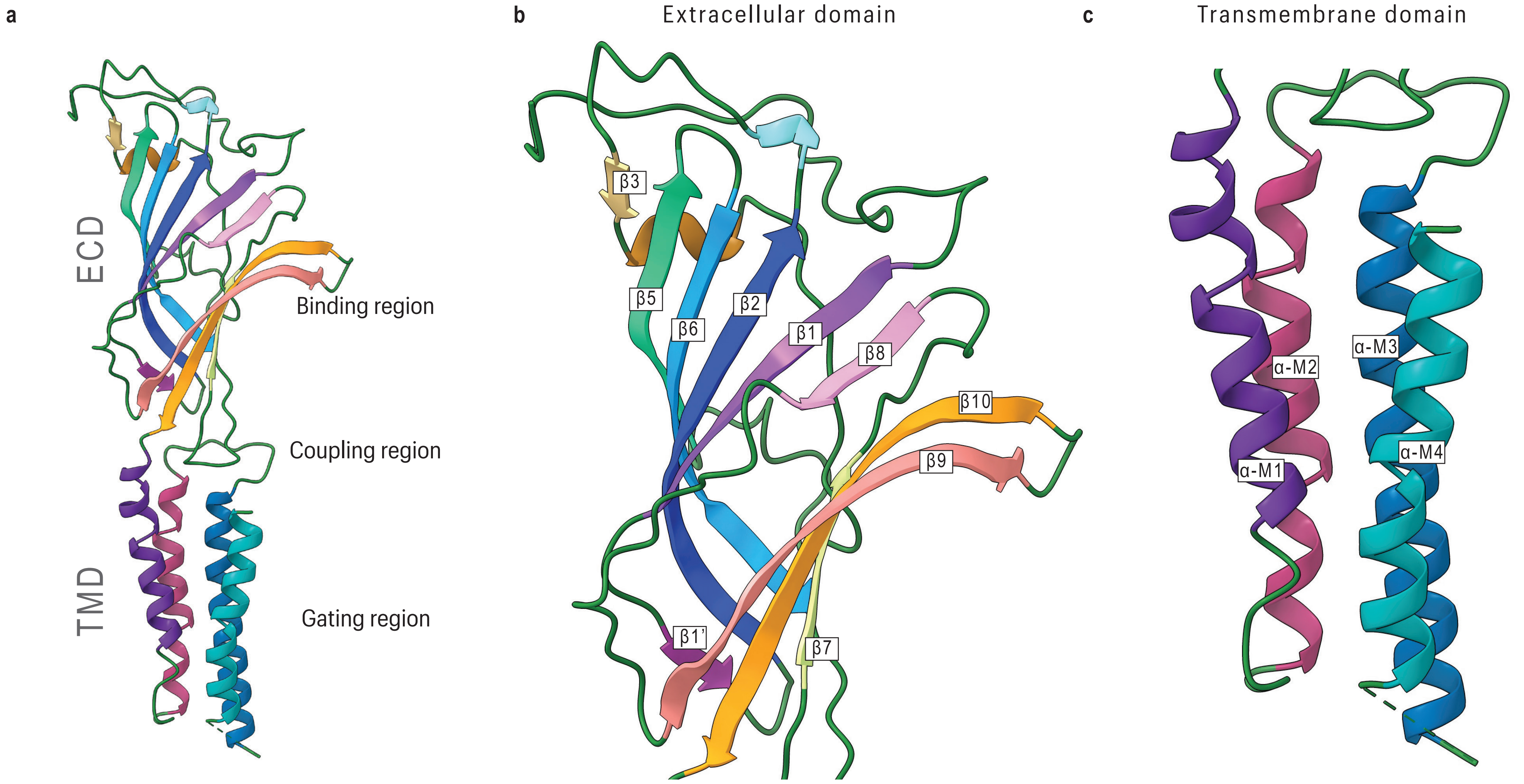

Supplementary Figure 12 -

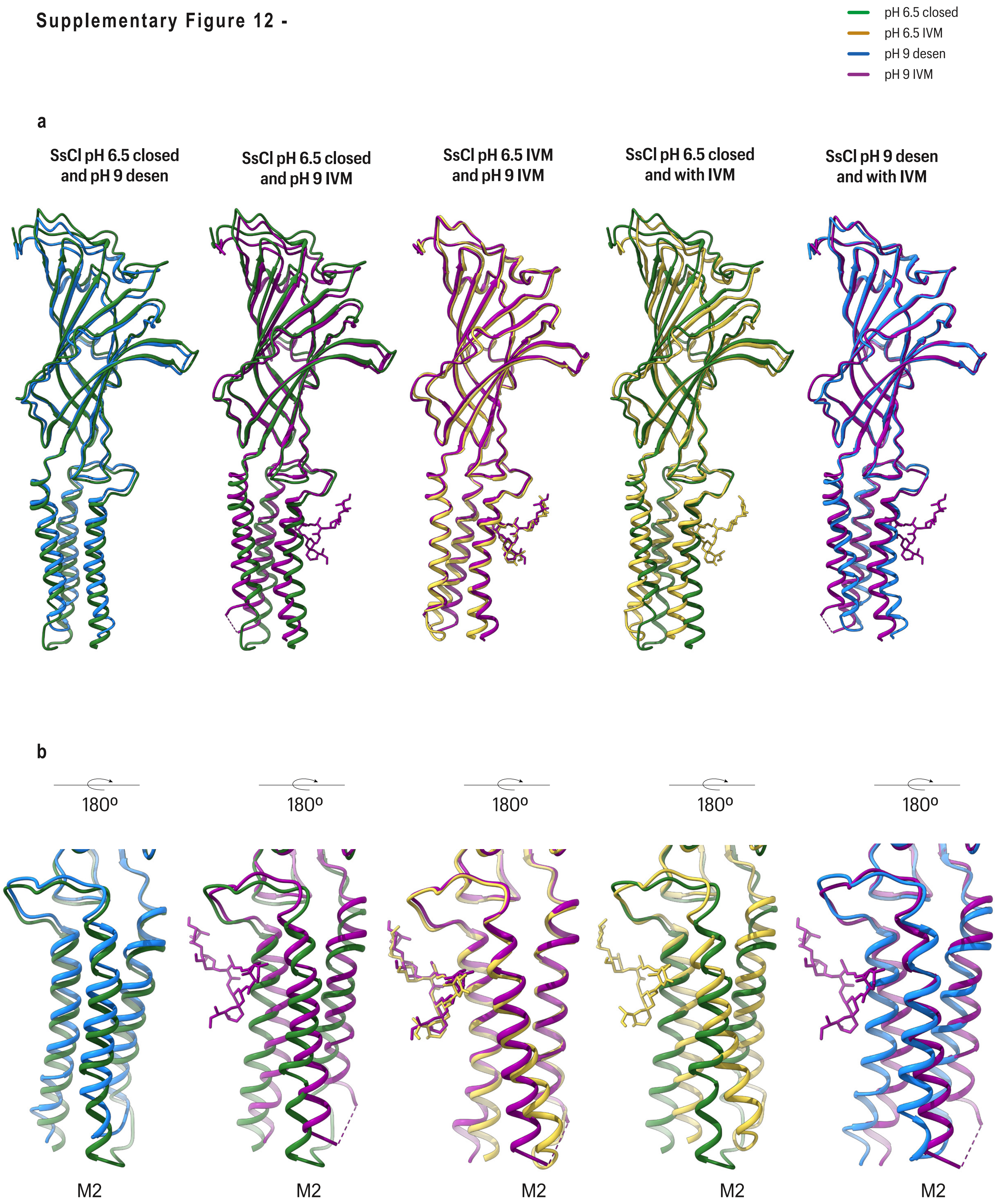

Supplementary Figure 13 -

SsCl pH 6.5 and pH 9 with IVM superimposed to GluCl bound to IVM

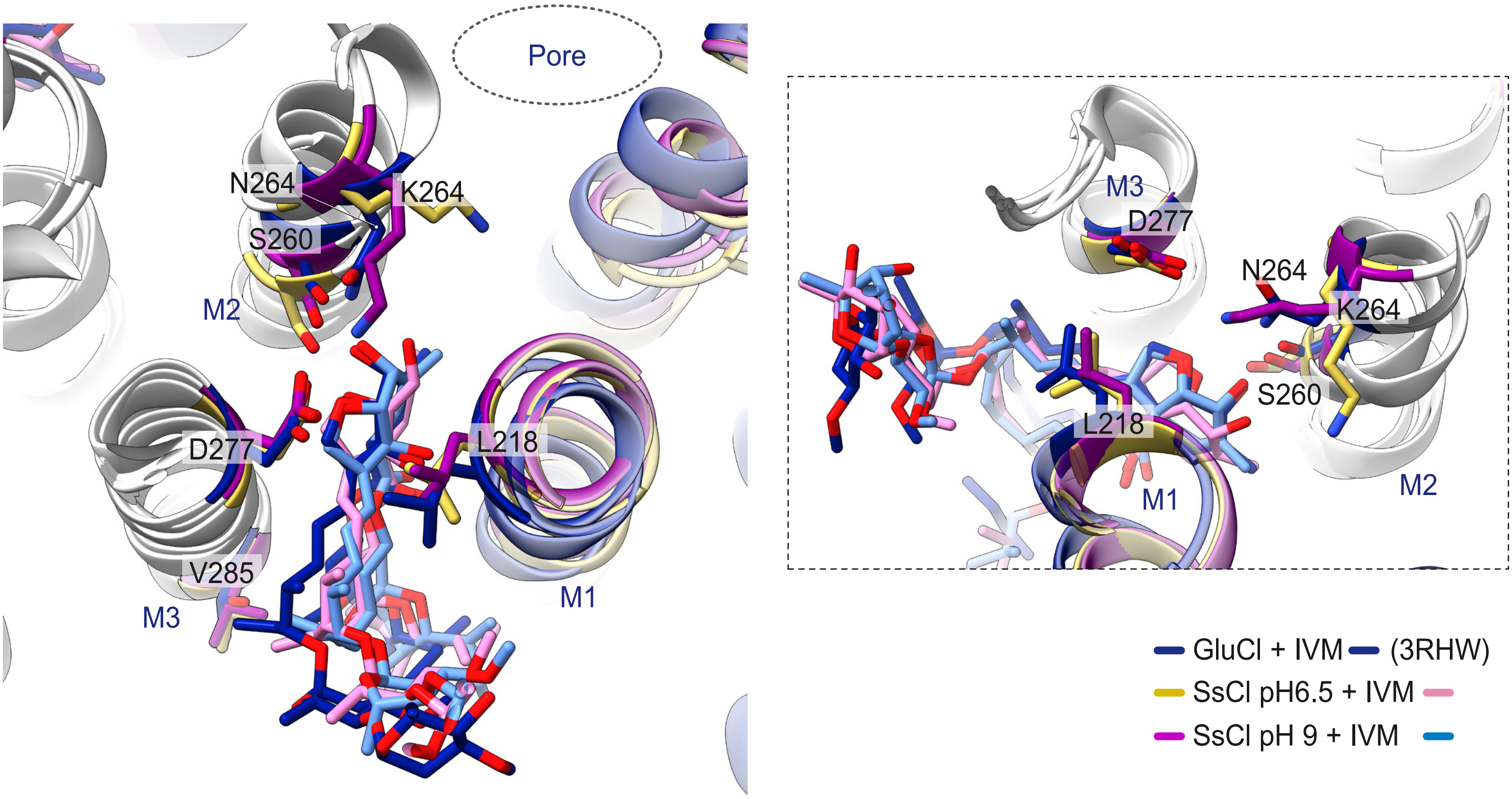

**Supplementary Figure 14 -**

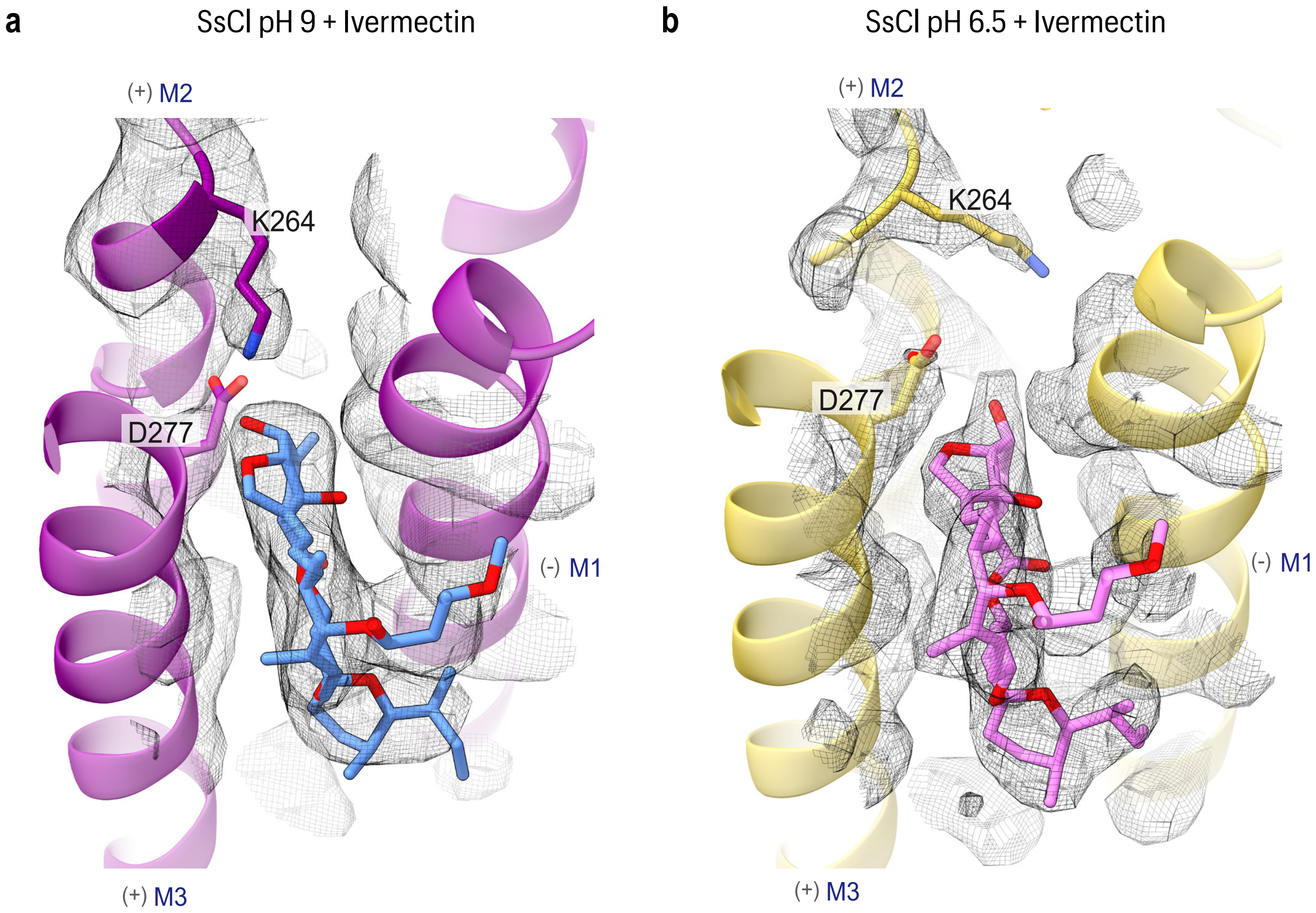

Supplementary Figure 15 -

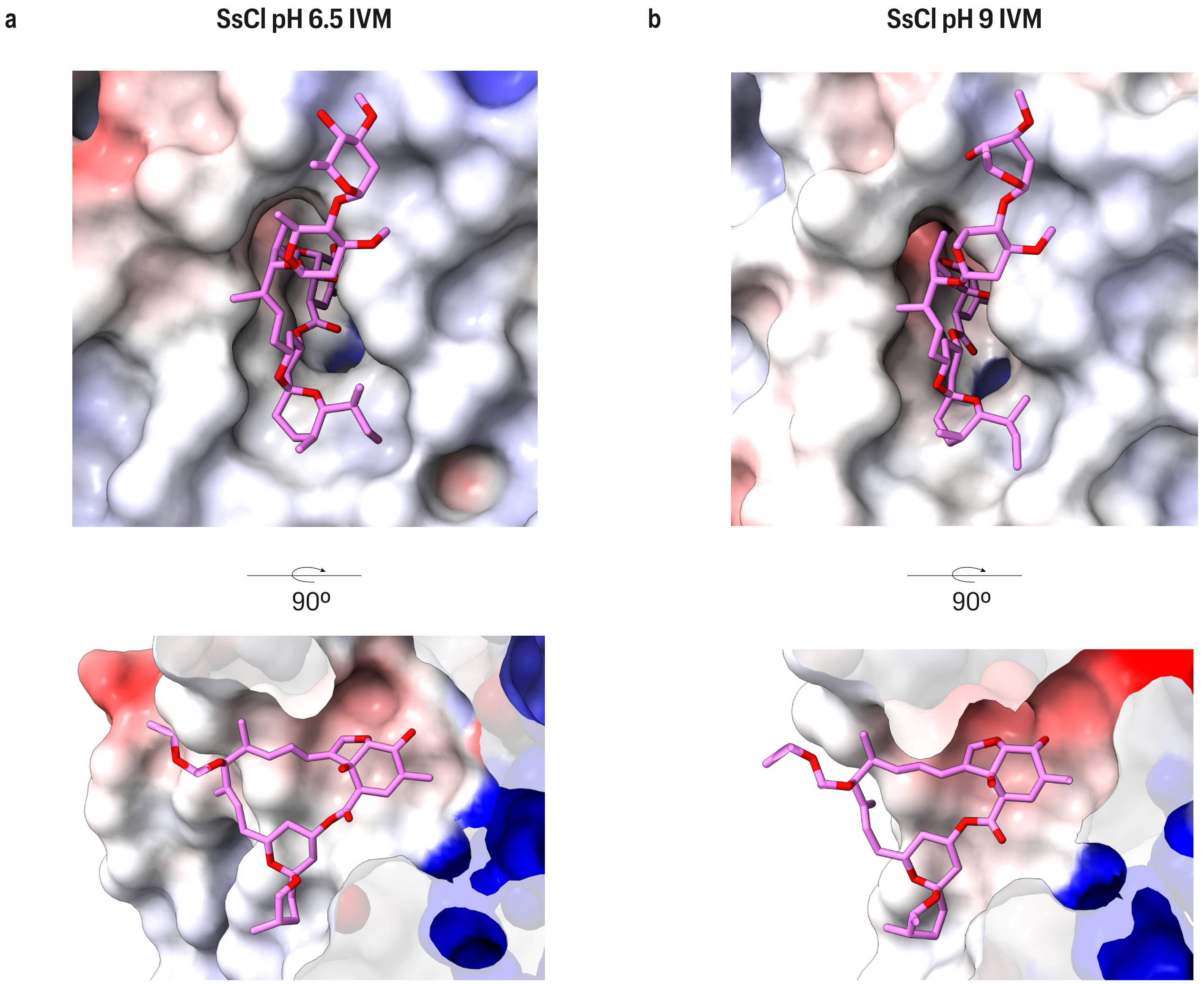

Supplementary Figure 16 -

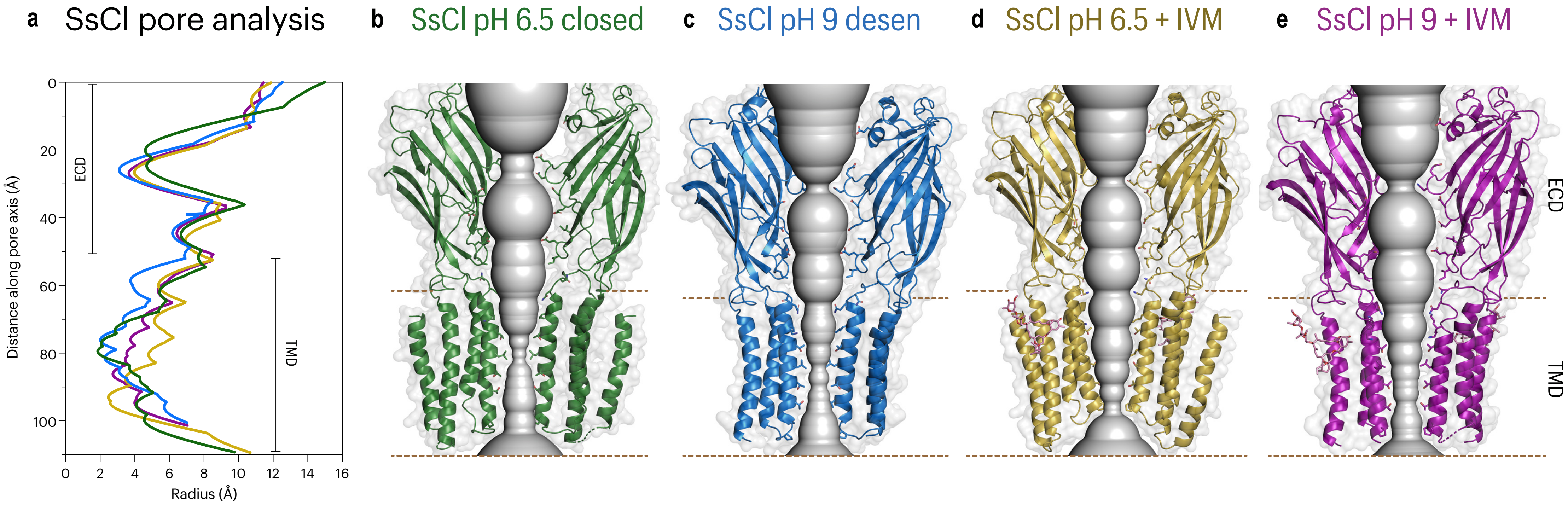

Supplementary Figure 17 -

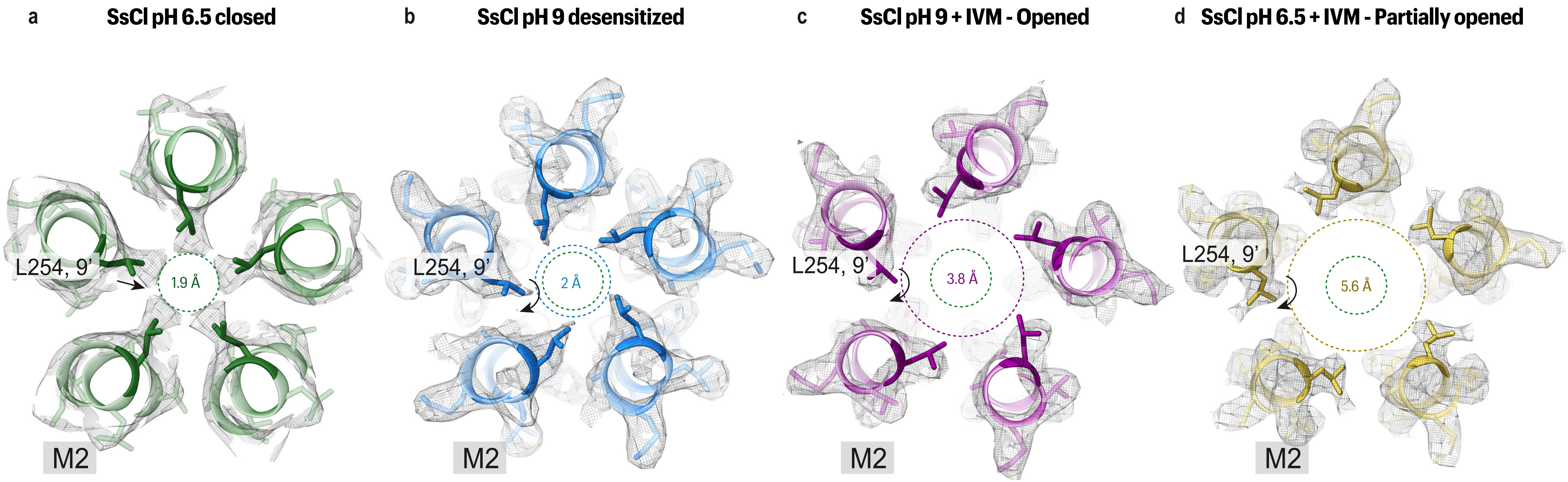

**Supplementary Figure 18 -**

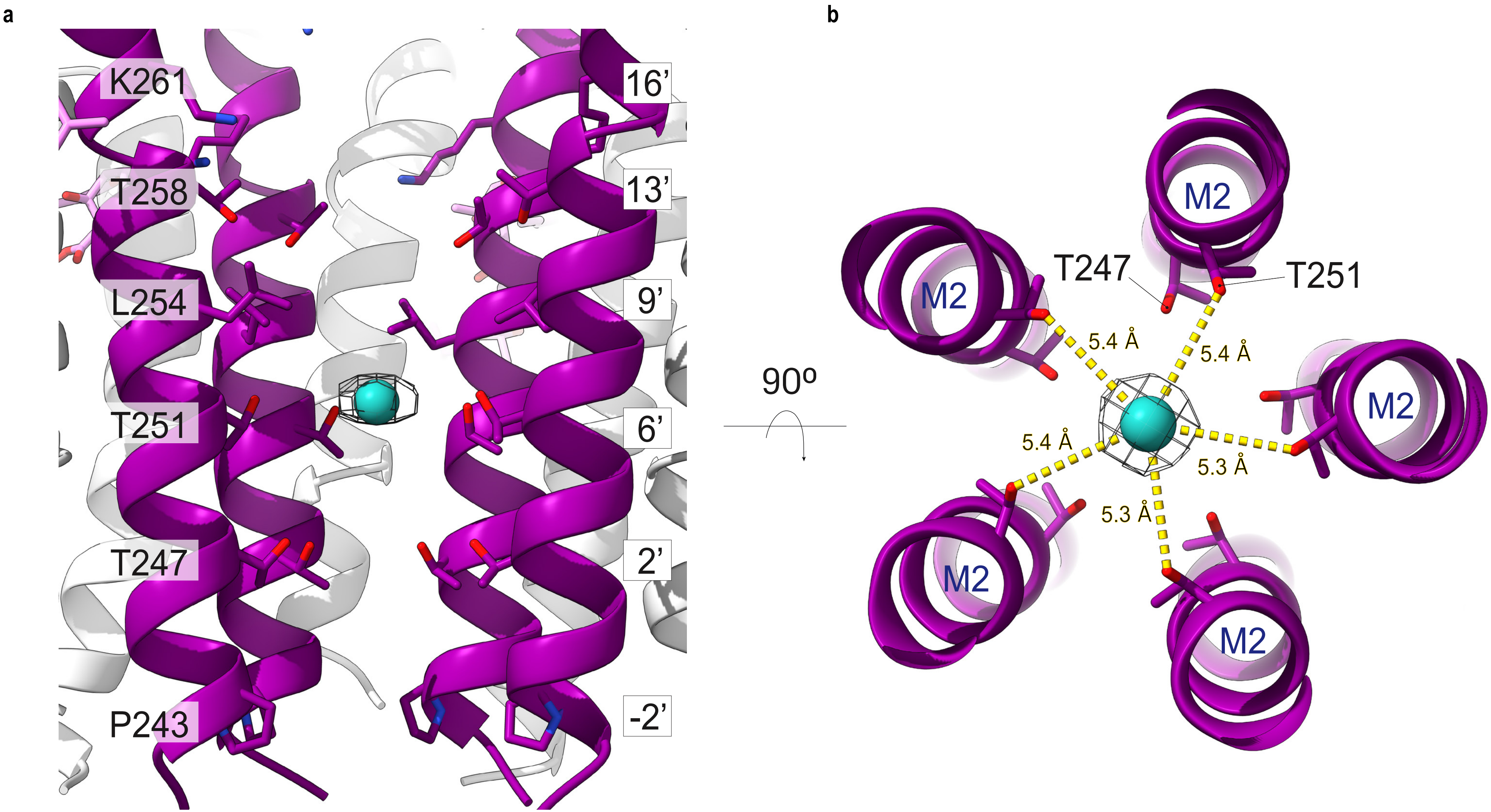

**Supplementary Figure 19 -**

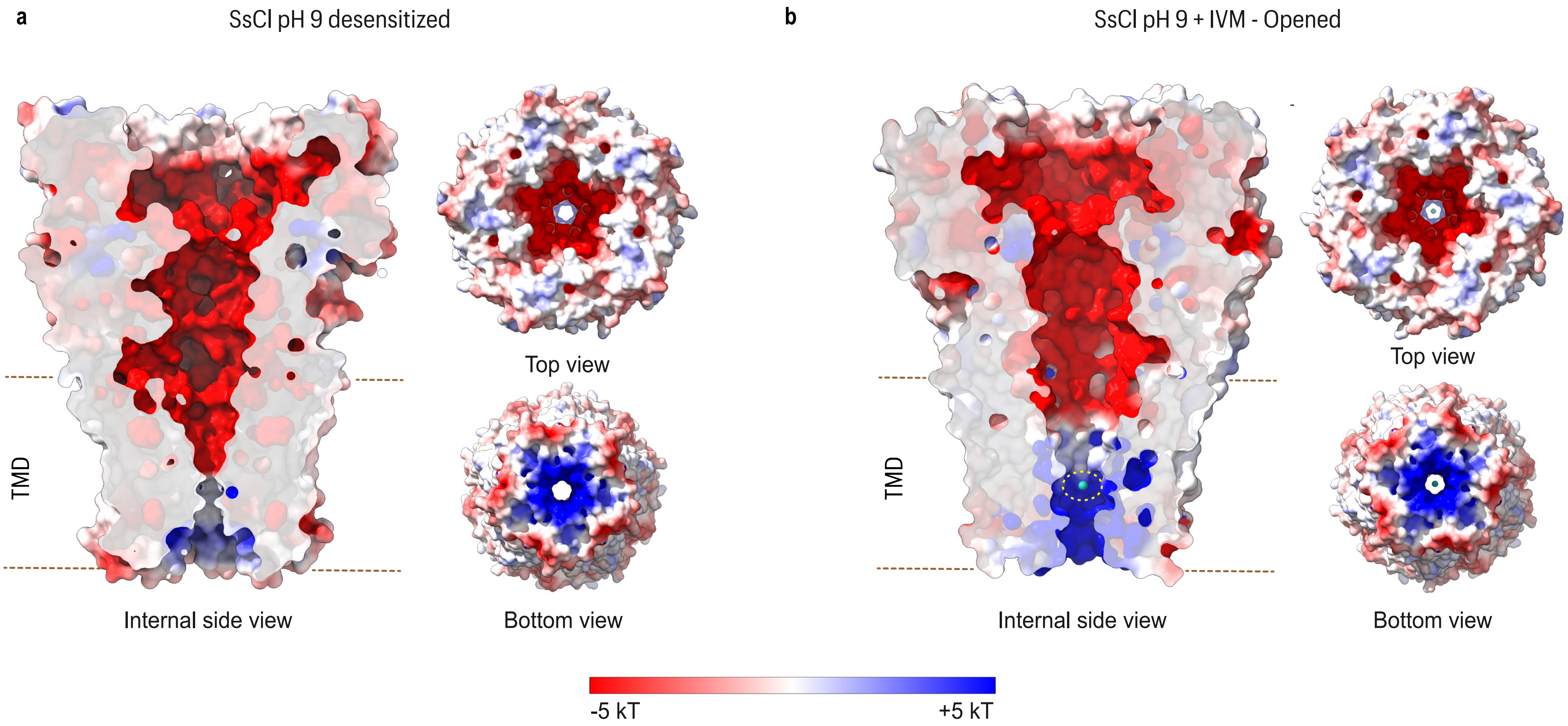

Supplementary Figure 20 -

a SsCl pH 9 desensitized

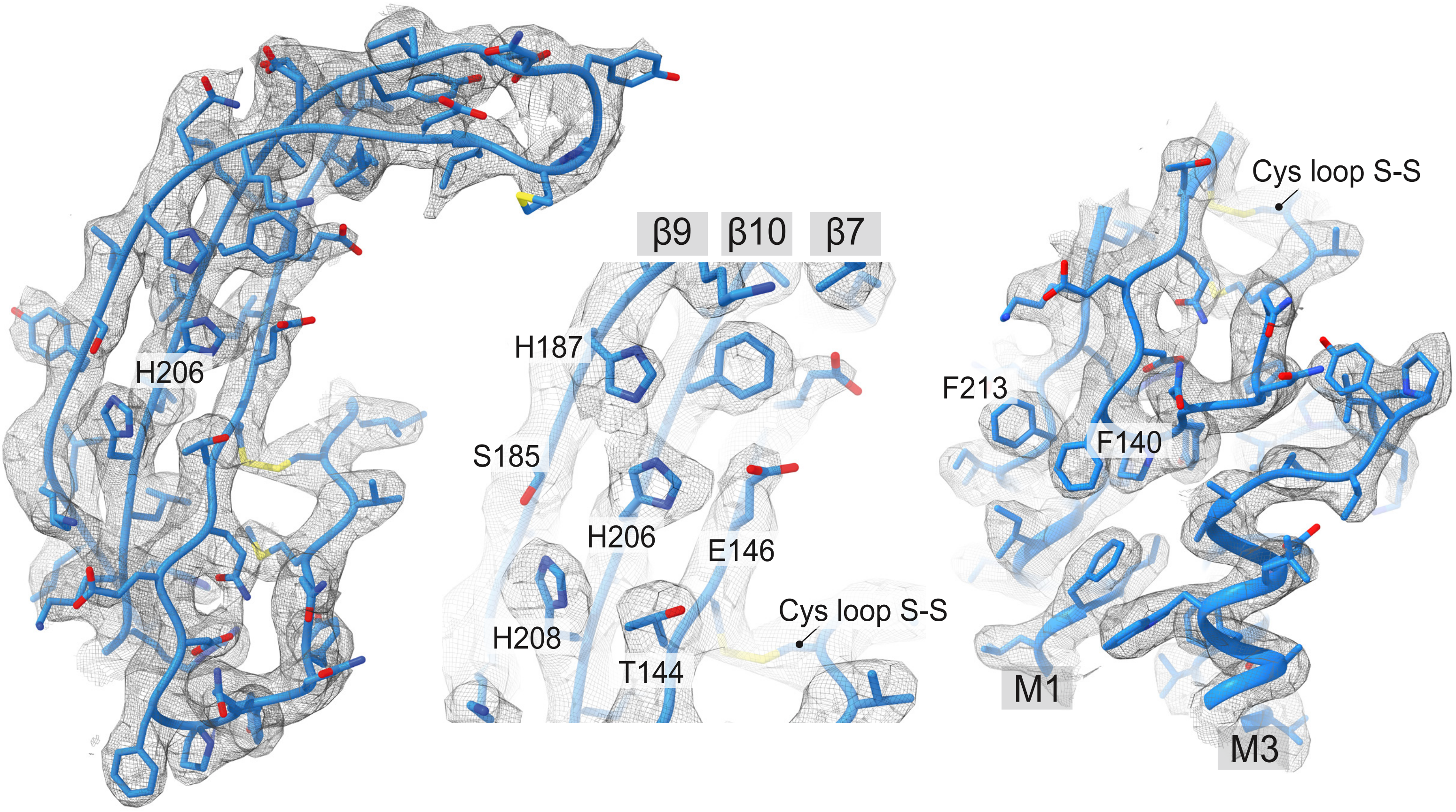


b SsCl pH 6.5 closed

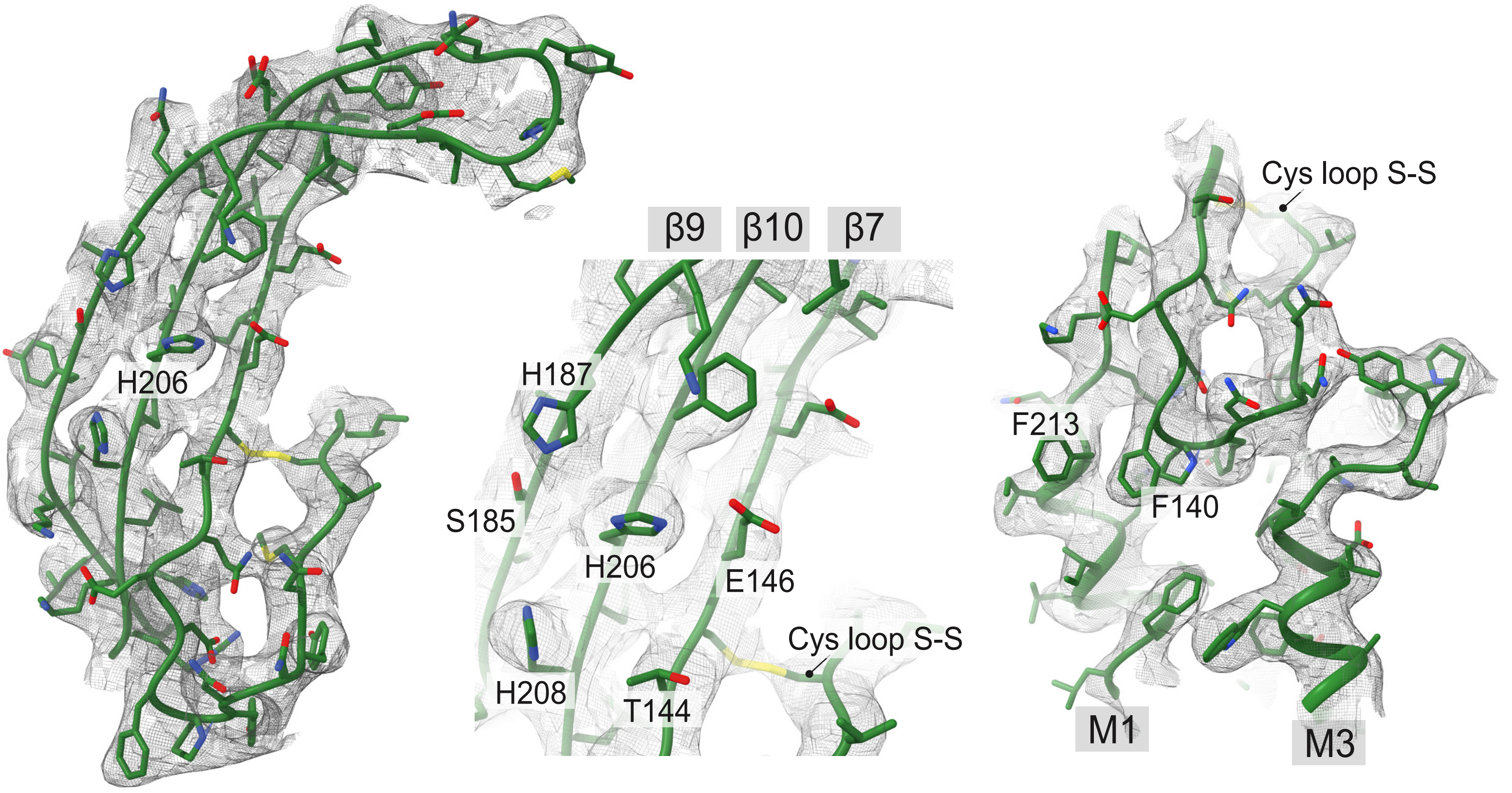

**Supplementary Figure 21 -**

**a**

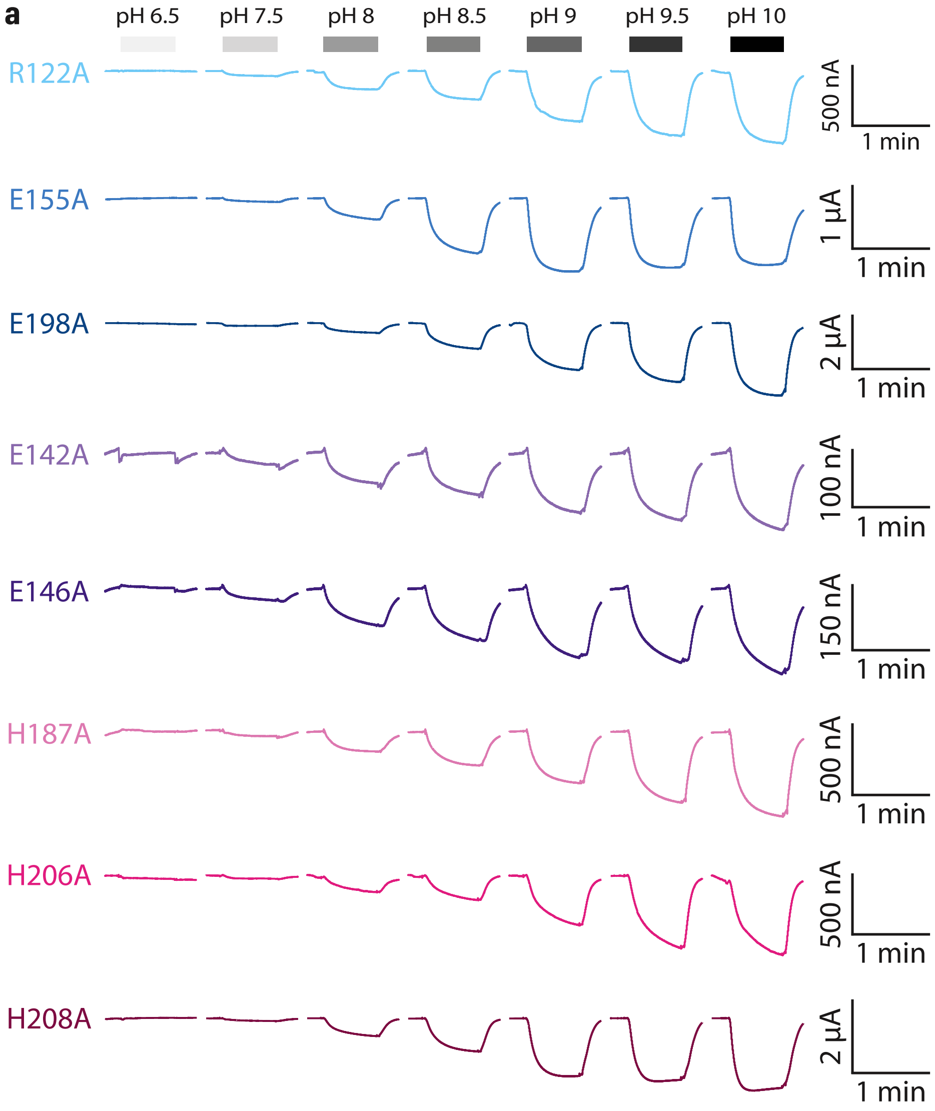


**b** [pH] Concentration responses

| Receptor | Max (nA) | nH | $pH_{50}$ | n |
|---|---|---|---|---|
| $SsCl_{EM}$ | 894.7 ± 145.7 | 1.12 ± 0.1 | 8.57 ± 0.04 | 4 |
| R122A | 313.0 ± 106.8 | 0.70 ± 0.04 | 8.82 ± 0.03 | 4 |
| E155A | 1208.0 ± 130.1 | 1.00 ± 0.13 | 8.33 ± 0.06 | 3 |
| E198A | 1480.3 ± 676.9 | 1.07 ± 0.14 | 8.51 ± 0.06 | 3 |
| E142A | 298.9 ± 80.4 | 0.74 ± 0.07 | 8.75 ± 0.06 | 3 |
| E146A | 268.8 ± 77.5 | 0.74 ± 0.03 | 8.92 ± 0.02 | 3 |
| H187A | 483.6 ± 47.7 | 1.14 ± 0.06 | 8.34 ± 0.02 | 3 |
| H206A | 572.3 ± 172.9 | 0.77 ± 0.06 | 8.88 ± 0.05 | 3 |
| H208A | 1475.7 ± 243.1 | 1.19 ± 0.06 | 8.46 ± 0.02 | 3 |
| H187A/H206A | 357.7 ± 77.6 | 0.70 ± 0.13 | 8.86 ± 0.33 | 3 |

**c** Application of IVM with pH 9

| Receptor | pH 9 | pH 9 + IVM | Rebound | n |
|---|---|---|---|---|
| Uninjected | -11.99 ± 0.99 | -0.01 ± 0.31 | 1.04 ± 0.29 | 5 |
| $SsCl_{EM}$ | 723.26 ± 78.38 | 3240.20 ± 316.49 | 5139.30 ± 482.72 | 10 |
| H206C/H187C | 95.04 ± 18.49 | 1039.03 ± 134.09 | 2003.90 ± 219.91 | 3 |

Supplementary Figure 22 -

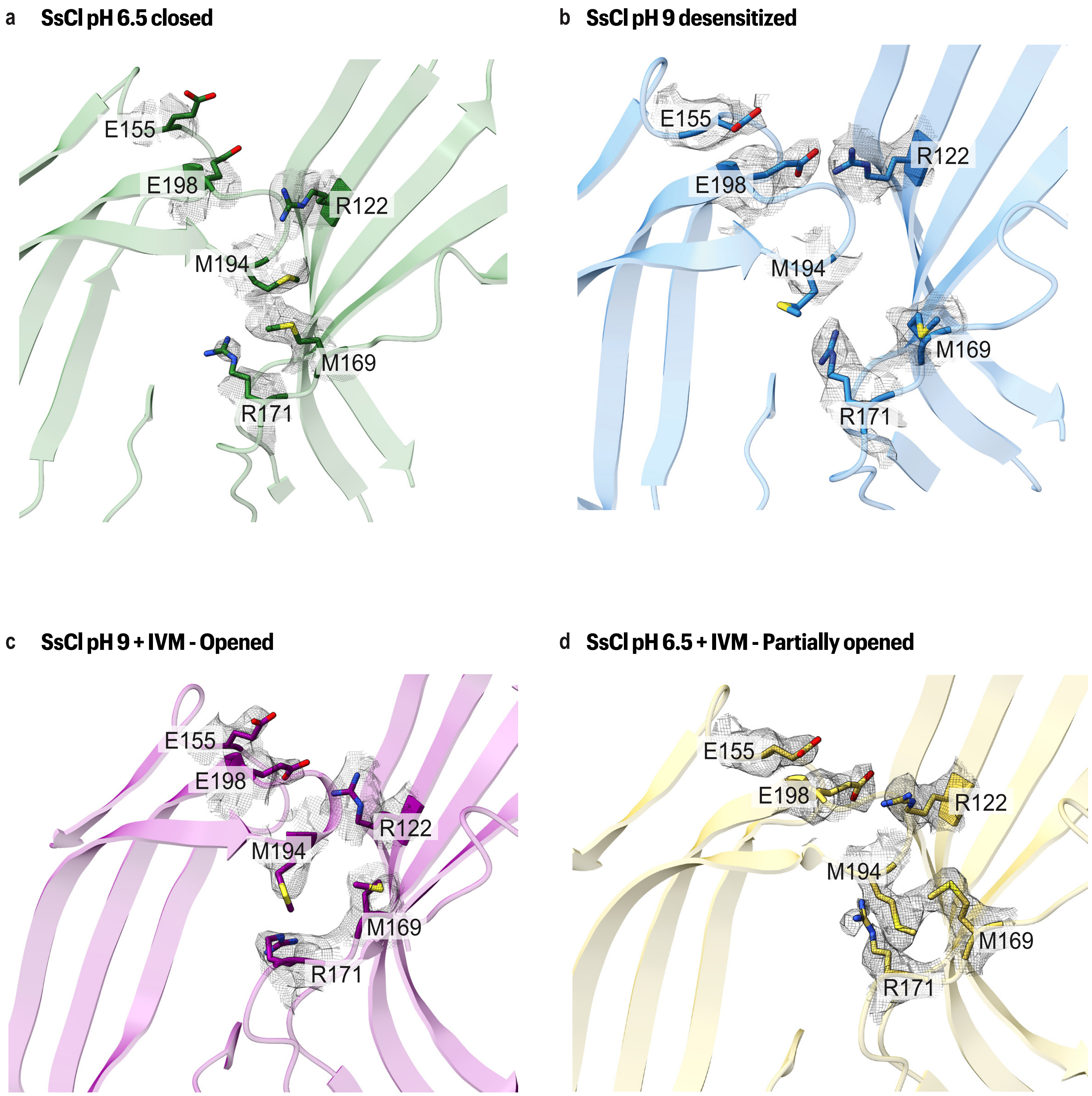

Supplementary Figure 23 -

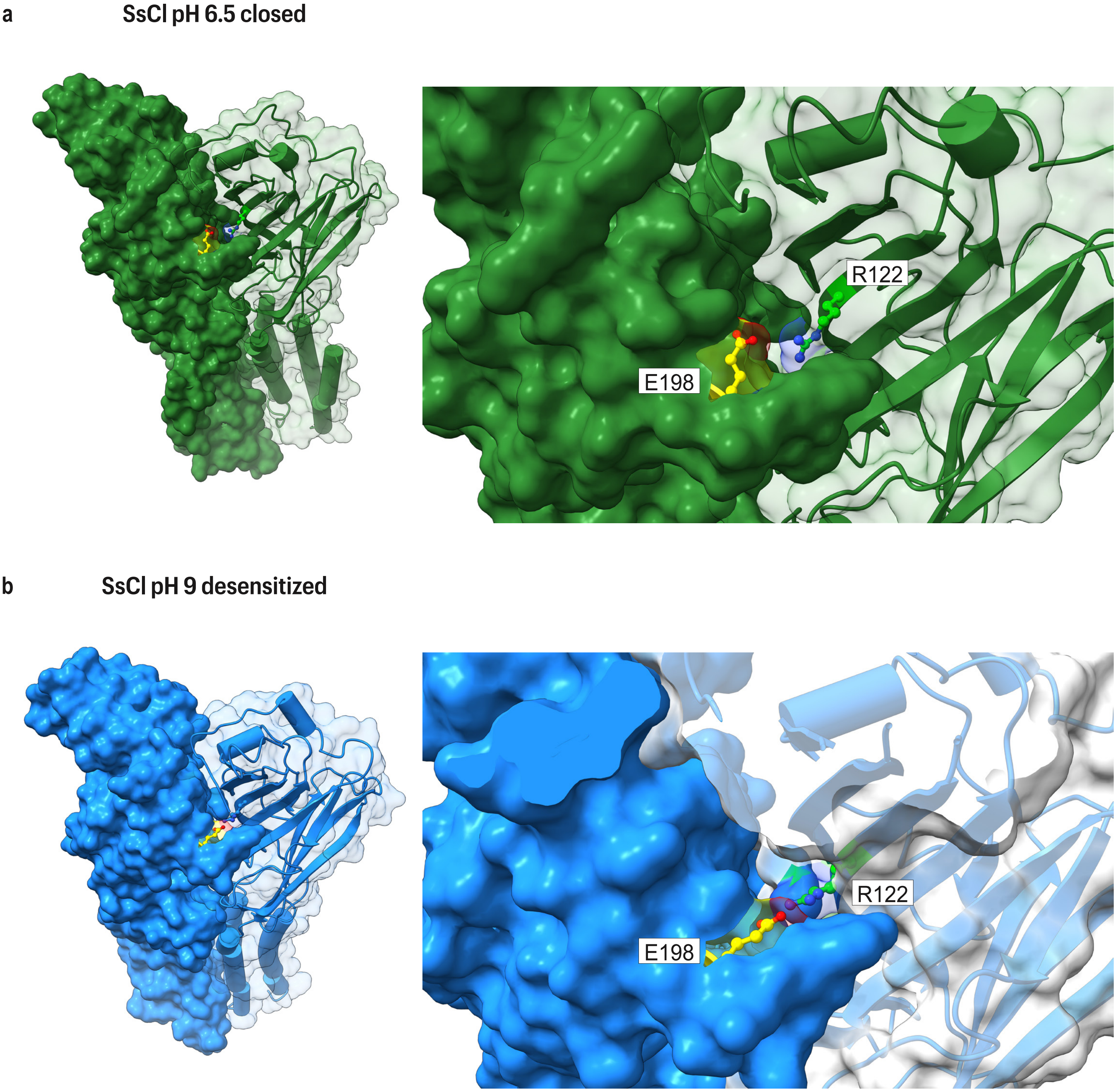

**Supplementary Figure 24 -**

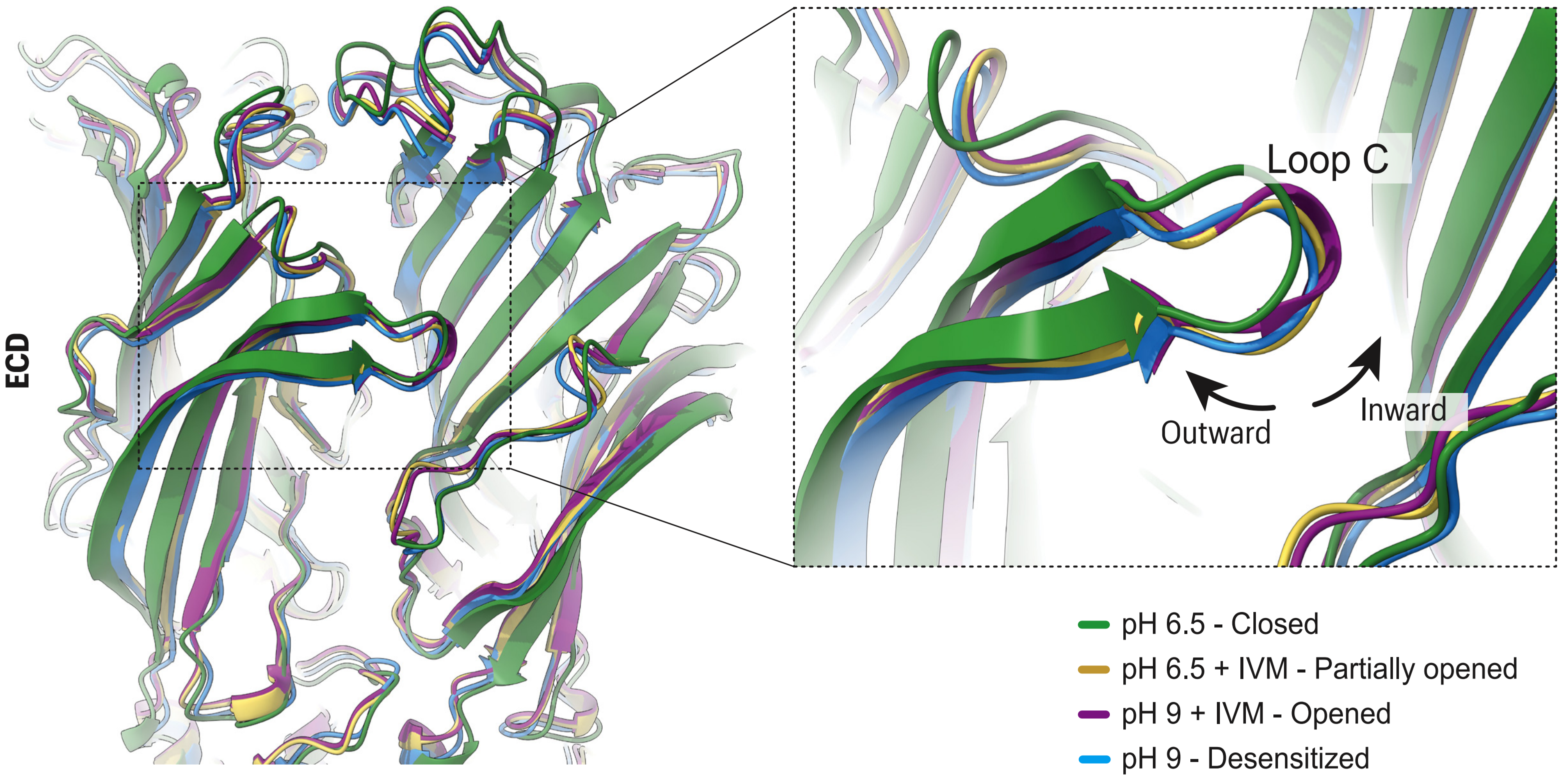

### Supplementary Table 1 - Cryo-EM info table

| | SsCl pH 6.5 closed<br>EMDB-53950<br>PDB 9RGM | SsCl pH 9 desensitized<br>EMDB-53952<br>PDB 9RGO | SsCl pH 6.5 + IVM<br>EMDB-53951<br>PDB 9RGN | SsCl pH 9 + IVM<br>EMDB-53953<br>PDB 9RGP |
|---|---|---|---|---|
| **Data collection and processing** | | | | |
| Microscope | Jeol CRYO ARM | Titan Krios | Titan Krios | Titan Krios |
| Magnification | 61,000x | 105,000x | 105,000x | 105,000x |
| Voltage (kV) | 300 | 300 | 300 | 300 |
| Electron exposure (e-/ Å 2 ) | 61 | 50 | 50 | 50 |
| Defocus range (μm) | 1.1 to 1.9 (0.2 steps) | -0.5 to -2.5 (0.3 steps ) | -0.5 to -2.5 (0.3 steps ) | -0.5 to -2.5 (0.3 steps ) |
| Pixel size (Å) | 0.720 | 0.831 | 0.829 | 0.829 |
| Symmetry imposed | C5 | C5 | C5 | C5 |
| Number of images | 7,366 | 23,382 | 50,059 | 29,306 |
| Particles picked * | 892,545 | 7,779,179 | 5,506,940 | 10,143,236 |
| Particles refined | 16,607 | 186,182 | 344,703 | 87,619 |
| Map resolution (Å) | 4.24 | 3.14 | 3.10 | 3.65 |
| **Model Building** | | | | |
| **Initial model used** | SsCl pH 9 desensitized | AlphaFold model | SsCl pH 6.5 closed | SsCl pH 9 desensitized |
| **Model refinement** | | | | |
| Model resolution (masked FSC 0.5) (Å) | 4 | 3 | 3 | 4 |
| FSCavg model mask FSC 0.5 (score) | 0.844 | 0.839 | 0.845 | 0.842 |
| Map sharpening *B* -factor (Å 2 ) (unmasked) | -168.57 | -121.12 | -108.34 | -153.25 |
| *B* -factor protein (Å 2 ) | 198.69 | 157.93 | 126.49 | 185.73 |
| *B* -factor ligand (Å 2 ) | None | None | 116.18 | 189.34 |
| **Model composition** | | | | |
| All atoms | 27,110 | 24,406 | 28,489 | 25,205 |
| H or D atoms (protein) | 13,480 | 12,135 | 13,863 | 12,210 |
| H or D atoms (other) | 0 | 0 | 370 | 370 |
| a.a. residues | 1,620 | 1,465 | 1,655 | 1,465 |
| Chains | 5 | 5 | 5 | 5 |
| Water | 0 | 0 | 0 | 0 |
| Ions | None | Chloride | None | Chloride |
| Ligands | None | None | Ivermectin (5) | Ivermectin (5) |
| **Model Validation** | | | | |
| Molprobity score | 1.74 | 0.96 | 1.50 | 1.50 |
| Clashscore | 5.05 | 0.94 | 5.30 | 6.98 |
| R.M.S.D - Bond (Å) | 0.0099 | 0.0110 | 0.0112 | 0.0099 |
| R.M.S.D - Angles (°) | 1.79 | 1.60 | 1.72 | 1.69 |
| Rotamers outliers (%) | 1.24 | 0.14 | 0 | 1.07 |
| Rotamers allowed (%) | 9.45 | 4.68 | 6.77 | 4.86 |
| Rotamers favored (%) | 89.32 | 95.18 | 93.23 | 94.07 |
| C-beta deviations | 30 | 5 | 11 | 19 |
| Ramachandran outliers (%) | 0 | 0 | 0 | 0 |
| Ramachandran allowed (%) | 5.94 | 3.02 | 3.34 | 2.42 |
| Ramachandran favored (%) | 94.06 | 96.98 | 96.66 | 97.58 |

**Supplementary Table 2 -**

| | | SsCl pH 6.5 closed | | SsCl pH 9 desen | |
|---|---|---|---|---|---|
| Residue | Position | pKa | Buried (%) | pKa | Buried (%) |
| GLU | 142 | 2.43 | 0 | 2.44 | 0 |
| ***GLU** | 146 | 4.30 | 25 | 5.09 | 47 |
| HIS | 187 | 5.93 | 0 | 6.02 | 6 |
| ***HIS** | 206 | 6.7 | 7 | 5.57 | 24 |
| HIS | 208 | 6.0 | 0 | 5.85 | 8 |
| GLU | 155 | 4.62 | 27 | 5.67 | 40 |
| GLU | 198 | 4.16 | 35 | 3.25 | 56 |

**Structure of a pH-sensitive pentameric ligand-gated ion channel from the *Sarcoptes* scabies mite**

Jessica Kleiz-Ferreira[1], Marijke Brams[1], Peter J. Harrison[2], Casey I. Gallagher[1], Mieke Nys[1], Ysaline Donze[3], Andrew Quigley[2], Daniel Bertrand[3], Chris Ulens[1].

[1]Laboratory of Structural Neurobiology, Department of Cellular and Molecular Medicine, Faculty of Medicine, KU Leuven, 3000 Leuven, Belgium.

[2]Membrane Protein Laboratory, Diamond Light Source, Research Complex at Harwell, OX11 0DE Didcot, United Kingdom.

[3]HiQscreen, 1222 Vésenaz, Geneva, Switzerland

### Supplementary Figure legends

**Supplementary Figure 1 | Sequence alignment of SsCl and other members of the pLGIC family.** Protein sequences were retrieved from public databases (NCBI) or internal resources. Secondary structure elements are annotated based on the SsCl reference sequence: β-sheets are shown in purple, β-sheet extensions in yellow, and α-helices in green. Species codes are: *Sarcoptes scabiei* (Ss), *Drosophila bipectinata* (Dbip), *Bactrocera oleae* (Bole), *Prorops nasuta* (Pnas), *Anabrus simplex* (Asim), *Bombyx mori* (Bmor), *Drosophila melanogaster* (Dmel), *Caenorhabditis elegans* (Cele), *Alvinella pompejana* (Apom), *Homo sapiens* (Hsap), and *Gloeobacter violaceus* (Gvio). The alignment was generated using the Clustal algorithm within the Jalview software, which was also used for visualization.

**Supplementary Figure 2 | Evolutionary analysis by Maximum Likelihood method of SsCl and other members of the pLGIC family.** Sequence alignment was generated using the Clustal algorithm within the Jalview software. Evolutionary analyses were conducted in MEGA11 [1]. The evolutionary history was inferred by using the Maximum Likelihood method and Le_Gascuel_2008 model [2]. The tree with the highest log likelihood -6649.91) is shown. The percentage of trees in which the associated taxa clustered together is shown next to the branches. Initial tree(s) for the heuristic search were obtained by applying the Neighbor-Joining method to a matrix of pairwise distances estimated using the JTT model. A discrete Gamma distribution was used to model evolutionary rate differences among sites (5 categories (+G, parameter = 2.6399)). The tree is drawn to scale, with branch lengths measured in the number of substitutions per site. This analysis involved 13 amino acid sequences. All positions containing gaps and missing data were eliminated (complete deletion option). There was a total of 265 positions in the final dataset.

Taxonomic classification (*Kingdom, Phylum, Class, and Order*) is shown for each species and color-coded accordingly. Species are: *Sarcoptes scabiei* (Ss), *Drosophila bipectinata* (Dbip), *Bactrocera oleae* (Bole), *Prorops nasuta* (Pnas), *Anabrus simplex* (Asim), *Bombyx mori* (Bmor), *Drosophila melanogaster* (Dmel), *Caenorhabditis elegans* (Cele), *Alvinella pompejana* (Apom), *Homo sapiens* (Hsap), and *Gloeobacter violaceus* (Gvio). PAC channel was used as outgroup.

Ref:
1. Tamura K., Stecher G., and Kumar S. (2021). MEGA 11: Molecular Evolutionary Genetics Analysis Version 11. Molecular Biology and Evolution https://doi.org/10.1093/molbev/msab120.
2. Le S.Q. and Gascuel O. (2008). An Improved General Amino Acid Replacement Matrix. Mol Biol Evol 25(7):1307-1320.

**Supplementary Figure 3 | SsCl wild-type and truncated constructs. a**, Sequence alignment is shown for the wild-type (SsCl WT) and truncated (SsCl$_{EM}$) constructs. Secondary structure elements are annotated based on the SsCl sequence: β-sheets (purple), β-sheet extensions (yellow), and α-helices (green). Key loop regions are also indicated. Cysteine residues involved in disulfide bonds are marked with a specific symbol (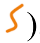). The intracellular domain truncation is indicated by a red line, with a scissor symbol denoting the truncation start site. The inserted linker sequence is highlighted in blue. Conserved residues between the SsCl WT and SsCl$_{EM}$ are marked with asterisks. **b**, Schematic representation of the constructs: SsCl WT with the full intracellular loop highlighted in red (left), and the truncated SsCl$_{EM}$ with the inserted linker shown in blue (right).

**Supplementary Figure 4 | Engineering, expression, and purification of truncated SsCl. a**, Schematic representation of the truncated SsCl$_{EM}$ construct engineered with an N-terminal GFP (green) for expression monitoring and fluorescence-detection size-exclusion chromatography (FSEC). **b**, Fluorescence microscopy of S*f*9 insect cells expressing the GFP-tagged truncated SsCl. **c**, FSEC profile of the GFP-tagged truncated SsCl, showing an oligomeric elution peak at ~16 mL (red). **d**, Schematic representation of the SsCl$_{EM}$ construct engineered with an N-terminal maltose-binding protein (MBP; yellow) for purification.

**Supplementary Figure 5 | pH sensitivity of wild-type and truncated SsCl in *Xenopus* oocytes. a,** Representative traces from oocytes injected with RNA encoding WT SsCl, SsCl$_{EM}$ or uninjected control oocytes. Pulses of pH 9 were applied in triplicate to assess pH-dependent activation. **b,** Representative trace showing concentration-dependent activation in an oocyte expressing SsCl$_{EM}$. **c,** Concentration-response curve for SsCl$_{EM}$. Data are presented as mean ± SEM ($n \geq 3$) and fitted using a modified Hill equation with a variable-slope linear regression model.

**Supplementary Figure 6 | Cryo-EM data processing workflow for SsCl.** Cryo-EM image processing pipelines for four SsCl datasets: **a**, SsCl at pH 6.5 (closed state); **b**, SsCl at pH 6.5 with IVM; **c**, SsCl at pH 9 (desensitized state); and **d**, SsCl at pH 9 with IVM.

**Supplementary Figure 7 | Local resolution estimation and global resolution assessment of SsCl cryo-EM maps.** Local resolution estimations for SsCl cryo-EM density maps were calculated using CryoSPARC and visualized in UCSF ChimeraX (right panels). Maps are colored according to local resolution, ranging from higher (blue) to lower (red) resolution. Global resolution was assessed by Fourier shell correlation (FSC), computed using the EMDB web server and visualized using the output graph from the server. FSC plots include: EMDB FSC (blue), 3σ threshold (orange dot-trace), and half-bit threshold (pink dotted), all calculated without mask. Standard FSC criteria, 0.5 (black traces), 0.333 (black dots), and 0.143 (green dot-trace), are shown as vertical lines. Additionally, CryoSPARC FSC tight (CS FSC, pink trace), calculated with mask, was also included in the graph. The CS FSC tight is considered the reliable estimate of global resolution for model interpretation, while the unmasked FSC at 0.5 serves as an additional reference for validation. **a**, SsCl at pH 6.5 (closed state); **b**, SsCl at pH 6.5 with IVM; **c**, SsCl at pH 9 (desensitized state); **d**, SsCl at pH 9 with IVM.

**Supplementary Figure 8 | Cryo-EM density of key structural regions in SsCl.** Cryo-EM density for a single SsCl subunit is shown for each of the four conformational states. Densities are shown for loop C, loop F, and the transmembrane helices (αM1–M4 for pH 6.5, and αM1–M3 for pH 9). Figures were generated using UCSF ChimeraX. **a**, SsCl at pH 6.5 (closed state); **b**, SsCl at pH 6.5 with IVM; **c**, SsCl at pH 9 (desensitized state); **d**, SsCl at pH 9 with IVM.

**Supplementary Figure 9 | Cryo-EM density of ivermectin in SsCl.** Cryo-EM density corresponding to IVM is shown for each structure. **a**, SsCl at pH 6.5 with IVM and **b,** SsCl at pH 9 with IVM.

**Supplementary Figure 10 | Cryo-EM density maps of SsCl at pH 9 highlighting the M4 transmembrane helices.** Cutaway side views and bottom views of the cryo-EM density maps reveal the obscured, poorly resolved α-M4 transmembrane helices, which are highlighted in red. **a**, **b**, SsCl at pH 9 (desensitized state); **c**, **d**, SsCl at pH 9 with IVM.

**Supplementary Figure 11 | Secondary structural features of SsCl.** The SsCl structure at pH 6.5 was used to illustrate conserved secondary structural elements present across all four states. **a**, Single subunit highlighting the typically binding, coupling, and gating regions of Cys-loop receptors. **b**, Extracellular domain with labelled β-strands. **c**, Transmembrane domain with labelled α-helices. Figures were generated using UCSF ChimeraX.

**Supplementary Figure 12 | Comparison of a single SsCl subunit across all states.** Structural alignments were performed in PyMOL and visualized using UCSF ChimeraX. **a**, Overlay of single subunits from different SsCl states, illustrating structural differences across the whole subunit. **b**, Alignment of the transmembrane region alone, rotated 180° relative to panel a.

**Supplementary Figure 13 | Structural comparison of the ivermectin binding pocket in SsCl and GluCl.** Top views of SsCl structures at pH 6.5 (yellow and pink) and pH 9 (purple and blue) with IVM bound, overlaid with the GluCl structure (dark blue, PDB ID: 3RHW). Key residues are represented as sticks. Structures were aligned and visualized using UCSF ChimeraX.

**Supplementary Figure 14 | Ivermectin binding pocket density in SsCl structures highlighting K264.** Side-tilted views of the IVM binding pocket show the position and cryo-EM density of K264 in different conformational states. **a**, SsCl at pH 9 with IVM, showing K264 oriented toward the ligand and potentially interacting with it, with well-resolved density. **b**, SsCl at pH 6.5 with

IVM, where K264 is rotated away from the ligand, supported by clear side-chain density. Figures were generated using UCSF ChimeraX.

**Supplementary Figure 15 | Electrostatic surface of the ivermectin binding pocket in SsCl.** Electrostatic surface potentials were calculated using APBS at the corresponding pH of each structure and visualized in UCSF ChimeraX. Side views (top panels) and side cutaway views (bottom panels, rotated 90°) are shown. **a**, SsCl at pH 6.5 with IVM; **b**, SsCl at pH 9 with IVM.

**Supplementary Figure 16 | Pore architecture of SsCl across four conformational states. a**, Pore radius profiles in Å (x-axis) plotted as a function of longitudinal distance along the channel, computed using HOLE software. **b-e**, Structural representations of SsCl in different states with pore rendered as light grey spheres: **b**, pH 6.5 (closed); **c**, pH 9 (desensitized); **d**, pH 6.5 with IVM; **e**, pH 9 with IVM. Figures were generated using PyMOL.

**Supplementary Figure 17 | Key pore-lining residue at the 9′ position and associated cryo-EM density.** Top views of the α-M2 transmembrane helices in SsCl structures highlight the conserved 9′ leucine (L254), shown as sticks with its corresponding cryo-EM density. Black arrows indicate the clockwise displacement of L254 across states. Pore radius is annotated at the channel center for each conformation. A green dotted circle, representing the pore radius of the pH 6.5 closed state, is overlaid in all panels for reference. **a**, SsCl at pH 6.5 (closed); **b**, SsCl at pH 9 (desensitized); **c**, SsCl at pH 9 with IVM; **d**, SsCl at pH 6.5 with IVM. Figures were generated using UCSF ChimeraX.

**Supplementary Figure 18 | Ion selectivity and chloride coordination in the open SsCl channel at pH 9 with ivermectin. a**, Cutaway side view of the TMD displaying pore-lining residues as sticks and the chloride ion (blue) coordinated by T251 at the 6′ position. **b**, Top view of the 6′ region, showing the chloride ion located 5.4 Å from the hydroxyl oxygen of T251. Figures were generated using UCSF ChimeraX.

**Supplementary Figure 19 | Electrostatic potential of SsCl at pH 9.** Electrostatic surfaces were calculated using the APBS software at pH 9 and visualized in UCSF ChimeraX. **a**, SsCl in the desensitized state; **b**, SsCl in the IVM-bound open state. Side-cut views (left) reveal the pore interior. Top and bottom views (right) are also shown. Surface coloring represents electrostatic potential: red (negative), white (neutral), and blue (positive). The chloride ion binding site is indicated with a yellow dashed circle, with the ion positioned based on cryo-EM density.

**Supplementary Figure 20 | Cryo-EM density of the pH-sensor region in SsCl.** Key residues within and surrounding the proposed pH-sensor are shown as sticks with corresponding cryo-EM density. **a**, SsCl at pH 9 (desensitized state); **b**, SsCl at pH 6.5 (closed state). Figures were generated using UCSF ChimeraX.

**Supplementary Figure 21 | Functional data for SsCl mutants. a,** Representative traces showing concentration-dependent activation in oocytes expressing $SsCl_{EM}$ with single-point mutations. **b,** Table containing the average maximum current, hillslope value (*nH*) and $pH_{50}$ for all $SsCl_{EM}$ constructs. Values were calculated by fitting concentration-response results with a variable-slope linear regression model. c, Table containing the average values for pH 9 with IVM co-application experiments on uninjected cells, $SsCl_{EM}$ and the H206C/H187C double mutant. For panels **b** and **c**, all data are shown as mean ± SEM.

**Supplementary Figure 22 | Cryo-EM density for key residues near loop C in SsCl.** Important residues in proximity to loop C are shown as sticks with their corresponding cryo-EM density. **a**, SsCl at pH 6.5 (closed); **b**, SsCl at pH 9 (desensitized); **c**, SsCl at pH 9 with IVM; **d**, SsCl at pH 6.5 with IVM. Figures were generated using UCSF ChimeraX.

**Supplementary Figure 23 | Interaction between R122 and E198 in SsCl.** One SsCl subunit is shown as a surface, with the complementary subunit displayed as a transparent surface and its backbone in cartoon representation. Residues R122 (green) and E198 (yellow), located near loop C, are highlighted as sticks. **a**, SsCl at pH 6.5 (closed state); **b**, SsCl at pH 9 (desensitized state). Figures were generated using UCSF ChimeraX.

**Supplementary Figure 24 | Conformational changes in loop C across SsCl states.** Structural comparison of SsCl reveals movement of loop C associated with channel activation. Structures are shown for: pH 6.5 closed (green), pH 9 desensitized (blue), pH 6.5 with IVM (partially open, yellow), and pH 9 with IVM (open, purple). A black arrow indicates the direction of loop C displacement during gating transitions. Figures were generated using UCSF ChimeraX.

### Supplementary Table legends

**Supplementary Table 1 | Cryo-EM info table.** Specifications regarding data collections, data processing and model building, including initial models, model refinements and model compositions for all four structures; SsCl pH 6.5 closed, SsCl pH 9 desensitized, SsCl pH 6.5 + IVM, SsCl pH 9 + IVM.

**Supplementary Table 2 | PROPKA values for residues predicted to contribute to the pH-sensor.** Residues highlighted from PROKPA analysis predicted to contribute to the pH-sensor of SsCl based on their pKa values and solvent-accessibility. Residues predicted to constitute the principal pH-sensor and form a key pH-sensitive ionic interaction are marked in bold with an asterisk (*).
