## Supplementary movie legends for "Structure of a pH-sensitive pentameric ligand-gated ion channel from the *Sarcoptes* scabies mite"

**Supplementary Movie 1 | Rotational movement of M2 helices in the SsCl desensitized state.** Morph from the closed structure at pH 6.5 to the desensitized state at pH 9, showing a top view of the desensitization movement. Rotational changes of L254 at the 9′ and –2′ positions are highlighted. Structural alignments were performed using MatchMaker in ChimeraX, and all visualizations were generated in ChimeraX.

**Supplementary Movie 2 | Concerted compression of SsCl in the desensitized state.** Morph from the closed structure at pH 6.5 to the desensitized state at pH 9, illustrating global compression from the side view. Structural alignments were performed using MatchMaker in ChimeraX, and all visualizations were generated in ChimeraX.

**Supplementary Movie 3 | β7 strand movement in the pH sensor region of SsCl at pH 9.** Morph from the closed structure at pH 6.5 to pH 9 structures, showing movement of the β7 strand in the pH sensor region. The desensitized state is colored blue, and the IVM-bound open state is shown in purple. Structural alignments were performed using MatchMaker in ChimeraX, and all visualizations were generated in ChimeraX.

**Supplementary Movie 4 | Rotation of M1 and M2 helices in SsCl at pH 9.** Morphs from the closed structure at pH 6.5 to pH 9 structures, highlighting top view rotation of the M1 and M2 helices. The desensitized state is shown in blue, and the IVM-bound open state in purple. Structural alignments were performed using MatchMaker in ChimeraX, and all visualizations were generated in ChimeraX.

**Supplementary Movie 5 | Rigid-body rotation of the transmembrane domain in SsCl IVM-bound states**. Morphs from the closed structure at pH 6.5 to IVM-bound states at pH 6.5 (yellow) and pH 9 (purple), showing rigid-body rotation of the transmembrane domain. Structural alignments were performed using MatchMaker in ChimeraX, and all visualizations were generated in ChimeraX.

**Supplementary Movie 6 | β7 strand movement in the pH sensor region of SsCl IVM-bound state at pH 6.5.** Morph from the closed structure at pH 6.5 to the IVM-bound state at pH 6.5, illustrating movement of the β7 strand near the pH sensor region. Structural alignments were performed using MatchMaker in ChimeraX, and all visualizations were generated in ChimeraX.

**Supplementary Movie 7 | Global structural compaction of SsCl.** Morphs from the closed structure at pH 6.5 to three target states: desensitized at pH 9 (blue), IVM-bound open at pH 9 (purple), and IVM-bound partially open at pH 6.5 (yellow), showing global compaction relative to the closed conformation. Structural alignments were performed using MatchMaker in ChimeraX, and all visualizations were generated in ChimeraX.

**Supplementary Movie 8 | Loop F conformational changes across SsCl structures**. Morphs from the closed structure at pH 6.5 to three target states: desensitized at pH 9 (blue), IVM-bound open at pH 9 (purple), and IVM-bound partially open at pH 6.5 (yellow), showing conformational changes of Loop F during channel transitions. Primary and complementary subunits are shown; Loop F is highlighted. Structural alignments were performed using MatchMaker in ChimeraX, and all visualizations were generated in ChimeraX.
